# G-quadruplex recognition by DARPIns through epitope/paratope analogy

**DOI:** 10.1101/2022.06.13.495947

**Authors:** Tom Miclot, Emmanuelle Bignon, Alessio Terenzi, Stéphanie Grandemange, Giampaolo Barone, Antonio Monari

## Abstract

We investigated the mechanisms leading to the specific recognition of Guanine Guadruplex (G4) by DARPins peptides, which can lead to the design of G4s specific sensors. To this end we carried out all-atom molecular dynamic simulations to unravel the interactions between specific nucleic acids, including human-telomeric (h-telo), Bcl-2, and *c-Myc*, with different peptides, forming a DARPin/G4 complex. By comparing the sequences of DARPin with that of a peptide known for its high affinity for *c-Myc*, we show that the recognition cannot be ascribed to sequence similarity but, instead, depends on the complementarity between the three-dimensional arrangement of the molecular fragments involved: the α-helix/loops domain of DARPin and the G4 backbone. Our results reveal that DARPins tertiary structure presents a charged hollow region in which G4 can be hosted, thus the more complementary the structural shapes, the more stable the interaction.

## Introduction

In addition to the well-known double helical arrangement, the important biological role of non-canonical nucleic acid is nowadays widely recognized. Among the different non-canonical DNA or RNA structures, guanine quadruplexes (G4s) are highly studied and characterized [1–3]. From a chemical point of view, G4s are formed in guanine-rich nucleic acids, whose nucleobases develop primarily cooperative Hoogsteen-type hydrogen bonds. Because of the specificity of this interaction, DNA (or RNA) is then organized in a series of stacked quartets, whose macromolecular arrangement is further stabilized by a metal cation occupying the central channel. Three main topologies can be adopted by non-canonical G4 structures depending on the 5’-3’ orientation of the strands forming the G4s backbone: parallel, antiparallel and hybrid (Figure 1). In the parallel conformation, all the strands are oriented in the same way, in the antiparallel conformation the strands are inversely oriented two by two, while in the hybrid conformation only one strand is inversely oriented with respect to the other three [4].

**Figure 1.**
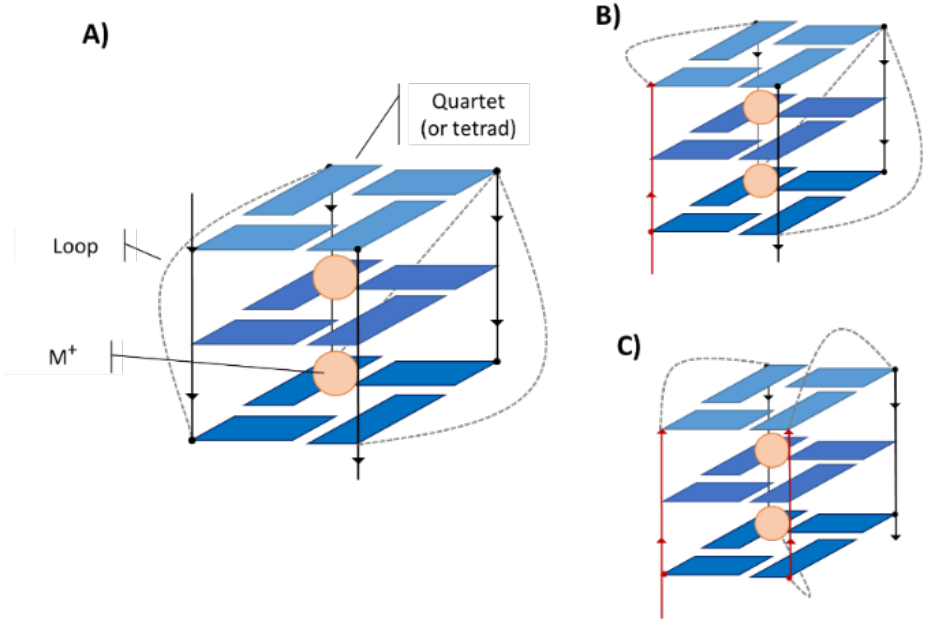
Schematic representation of the different topologies of G4. A) The parallel form in which all strands are parallel to each other, B) the hybrid form in which one strand is oriented antiparallel to all the others, and C) the antiparallel form in which the strands are antiparallel two by two.

G4s have been identified in cellular nucleic acids and have been associated to the control of key biological functions. As a matter of fact, G4s are involved in gene regulations [5,6], in neurodegenerative diseases [7,8], in the induction of DNA damages and in oncogenesis [9–12]. Furthermore, they have been recognized to play a role in the regulation of DNA cycles and in the regulation of post-translational modification in proteins [13– 16]. G4s have also been identified in both DNA and RNA viral genomes, including SARS-CoV-2 [17,18], where they may exert vital functions in regulating viral infection cycles [19–22]. Obviously, all these processes can only take place through a molecular machinery involving proteins selectively recognizing specific DNA or RNA G4s. Among them we may cite ATP-dependent DNA/RNA helicase DHX36 [23], G-rich sequence factor 1 [24], or fragile X mental retardation protein (SFMRP) [25]. Thus, the development of artificial or biomimetic specific G4 binders, capable to recognize either DNA or RNA, is highly valuable. Furthermore, such ligands may be exploited either in a therapeutic context or for the rapid identification and localization of G4s in cells or cellular compartments [26–29]. Protein engineering has also led to the development of antibodies presenting specificity and selectivity toward G4s. In this case, the recognition of G4s proceeds through the epitope/paratope mechanism, in which the G4 acts as the epitope of an antibody [30]. However, the design of antibodies is definitively not straightforward, and their use is typically limited to the identification of the subcellular localization of G4s [31–34]. Smaller peptides specifically recognizing G4s and even discriminating between different G4 types have also been proposed. This is tipycally the case of DARPins [35], a class of synthetic proteins derived from the modification of natural ankyrins and mostly known as chaperone agents in crystallography [36]. In addition, DARPins have also been used as cellular markers in biological imaging and for therapeutic purposes [37,38].

Understanding the factors underlying the specific recognition of G4s by DARPins can facilitate the design of sensors able to discriminate the G4s subcellular localization and their specific sequences. In this contribution, we model the DARPin/G4 interaction and, thus, unravel the specific recognition modes by combining molecular docking and long-scale all atom molecular dynamics (MD) simulations. We focus on 2E4 DARPin, which is specific for the G4 present in the *c-Myc* oncogene promoter, and 2G10, which has a slight specificity for different G4s [35]. As for the nucleic acid counterpart, we restrict our study to the human telomeric G4, as well as the G4s in the *c-Myc* [35] and *Bcl-2* promoters.

## Results and Discussion

MD simulations shed light on the structural details underlying the specific DARPins/G4 interaction. Indeed, by sampling the conformational space through different initial interaction positions, it is possible to analyze whether the interaction is conserved, the binding of the G4 affects the flexibility, the nucleic acid rearranges to reach a more stable pose, or if the proposed DARPin/DNA complex is not stable and separates. In our case most of the G4s/DARPin complexes are persistent and stable all along the MD and the peptides interact with the G4s through regions composed of large loops and helices, which overlap well with the recognized canonical interaction zones of the DARPins.

The only exceptions can be highlighted for 2E4/*c-Myc* which in one of the poses leads to a very labile and mobile interaction between G4 and the protein as confirmed by clustering yielding to two dominant structures, representing 41.71% and 30.56% of the trajectory, respectively. On the other hand,2E4/*h-Telo* (64.33% of thetrajectory) and 2G10/*c-Myc* (75.10%) yield dominant clusters that interacts only through the loop ends of DARPins and one or two nucleotides of the flexible G4s loops. (All the clustered structures can be found in the Supplementary Information).

### Residue-scale analysis of the G4/DARPin complexes

Before exposing the structural details of the G4s/DARPin complexes at the atomistic scale, it is interesting to consider the interaction at a residue-level scale, and in particular classify the different interaction patterns in terms of the number of involved nucleic acid or protein residues. Figure 2 show all the residues which remains within a cutoff of 3 Å from either the protein or the nucleic acid with a frequency at least equal to 50% of the simulation time. On average six nucleic acid residues of *h-Telo* and eight amino acids of 2G10 can be identified. However, 2G10 interacts persistently through only five amino acids with *c-Myc* and *Bcl-2* which in turn only bring a maximum of two or three nucleotides into persistent contact with the protein. Conversely, the 2E4/*h-Telo* interaction appears to be driven by three nucleotides and six amino acids. For 2E4, the interaction gathers eight amino acids with both *c-Myc* and *Bcl-2*, yet a different number of nucleic acid residues is involved, i.e. four for *Bcl-2* and seven for *c-Myc*.

**Figure 2.**
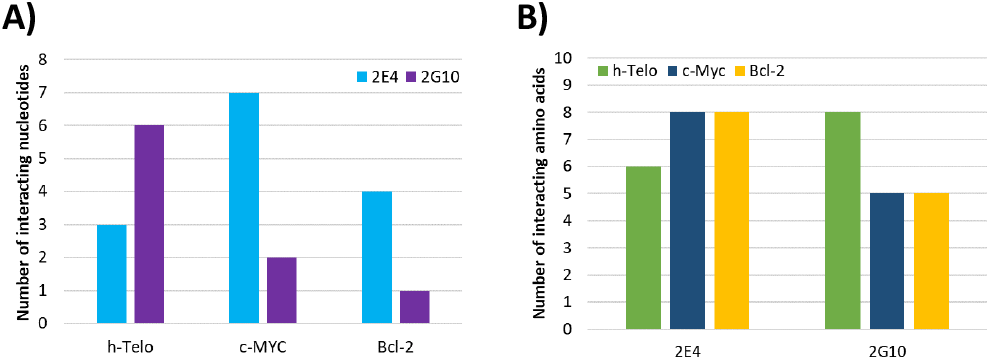
Number of nucleic acids (A) and amino acids (B) residues involved in the protein/DNA interaction and found at a frequency of at least 50% of the simulation time.

This first analysis, at the residue level, already draws a general picture of the specific recognition of G4 by DARPins, highlighting that the protein/nucleic acid recognition is favored by a high number of interacting residues. However, it needs to be completed identifying the exact nature of the interacting residues, their specific frequency, and the specific structural features.

### Three different modes of interaction leading to the G4/DARPin complexes

To improve the global scale analysis presented in the previous subsection we should identify a region of the G4s that is selectively recognized by DARPins. As a matter of fact, *h-Telo* does not show any specific interacting region or hotspot with 2E4 or 2G10 (see Figure 3A). This could confirm that the non-specific recognition of *h-Telo* is due to the absence of a well-defined target region on this nucleotide. In contrast, more pronounced specific interaction regions may be recognized in the two others G4s. Indeed, it can be seen in Figure 3B that two interaction areas clearly stand out for *Bcl-2* interacting with either 2EA or 2G10, i.e. the one including residues dC5 to dG8 and the one involving residues dG20 to dG22. Since the same nucleic acid regions are evidenced for both DARPins, the interaction mode can be classified as structurally similar in each complex. Thus, no specific recognition of *Bcl-2* by 2E4 or 2G10 can be inferred, since such specific recognition should involve interaction areas that must differ between two different DARPins. This is, indeed, the case for the G4 present in the *c-Myc* promoter. Figure 3C clearly shows regions of very pronounced contact and different for each of the DARPins. For the interaction with 2G10, the hot spot includes residues dG15 to dT20, although the contact frequencies are still quite spread across the whole G4. On the other hand, 2E4 highlights two very strong and localized contact points. The most important one concerns residues dG6 to dA12, while the second one corresponds to the last two residues of G4, dA21 and dA22. Thus, the specific recognition of *c-Myc* by 2E4 could be achieved either through the recognition of its sequence, or through a specific structural motif. Our analysis indicates three possible scenarios: 1) a rather general interaction that does not involve any specific G4 region or sequence (*h-telo*); 2) a non-specific interaction involving particular G4s regions, which are however recognized by all the proteins (*Bcl-2*); 3) a specific interaction driven by a few nucleotides having very high contact frequencies with specific DARPin (*c-Myc*).

**Figure 3.**
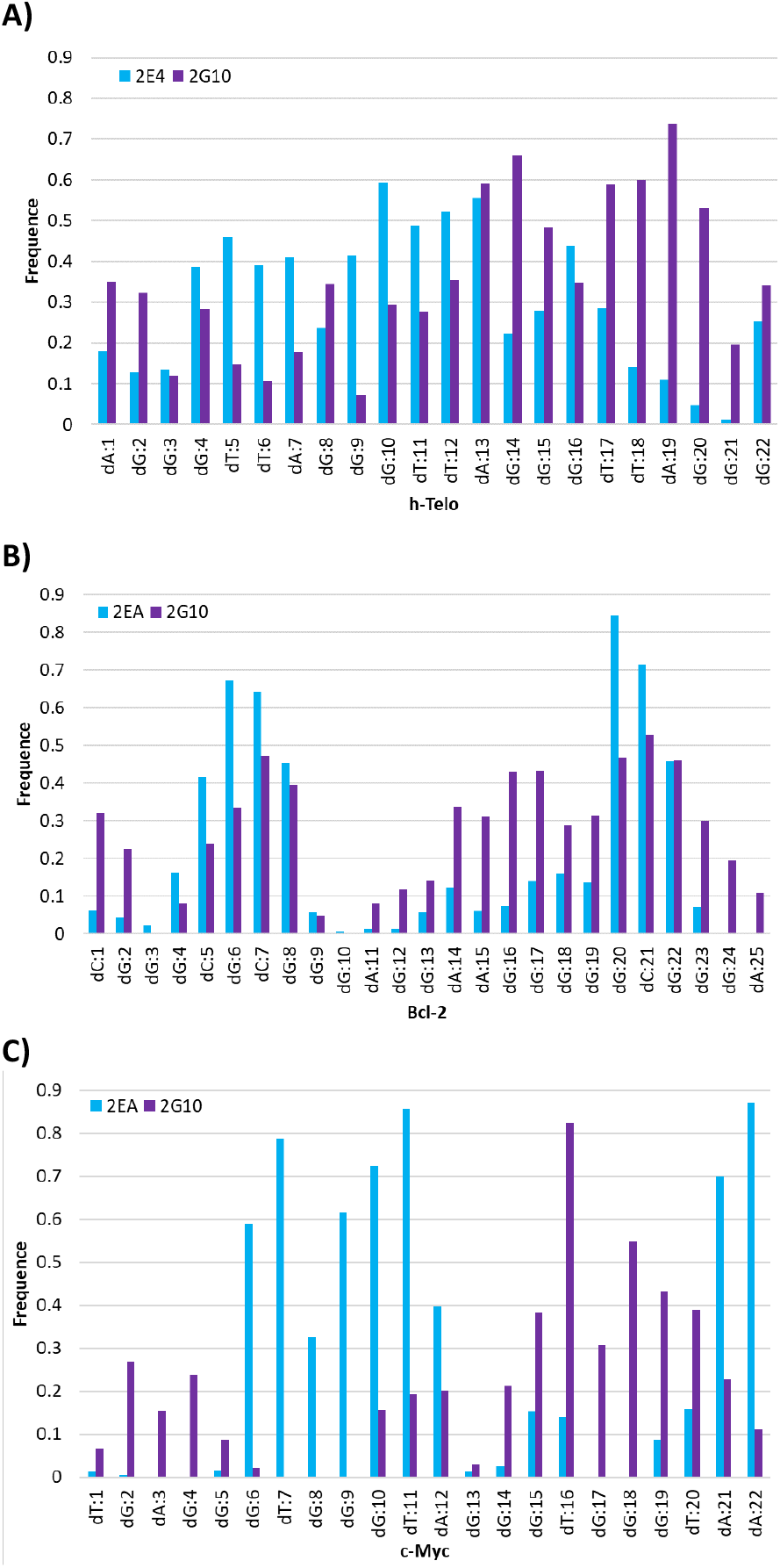
Nucleotide frequency of A) h-Telo, B) Bcl-2 and C) c-Myc involved in a 3 Å cutoff of the protein.

### Identification of a putative selective DARPin interaction area

Repeating the same analysis while focusing on the protein counterpart we identify the amino acids mostly involved in the recognition of the non-canonical DNA structure. Similarly, to what has already been observed for G4s, selectivity should correlate with few specific aminoacids having high contact frequencies with G4s. Conversely, for non-selective recognition a more scattered distribution of the interaction frequencies should be observed.

The distribution of the interaction contacts of 2G10 with the three G4s (Figure 4A) shows three distinct peaks. The first one corresponds to residues N34 and I35, the second one gathers residues R67, W68, R70, K78 and W79, while the last one comprises residues K100 and K101. While these localized protein areas certainly correspond to a strong interaction with G4, they appear rather non-specific since they are present for all the three G4s. However, the interaction with *h-Telo* is also driven by amino acids whose contact frequency was low or zero for the other G4S. This case concerns mainly residues H107, L108, I111, R112, K133, F134, K136, and I141. However, caution should be taken to avoid overinterpretation of this result, since 2G10 is not showing any specificity for *h-Telo* [35]. The contact frequency for 2E4 (Figure 4B) shows the emergence of even more defined trends presenting stronger and more localized maxima. In particular, we can mention residues K5, E9, and R12 as well as the regions spanning residues R34, W35, and M46, and residues H67, W68 and R70. However, only a relatively small difference in the interaction patterns between the three G4s can be highlighted. In particular in the case of *c-Myc* residues Y45, R70, L75, S78, R79, and G80 develop persistent contacts, and hence could be regarded as potential hot-spots for the selectivity of 2E4 towards this G4.

**Figure 4.**
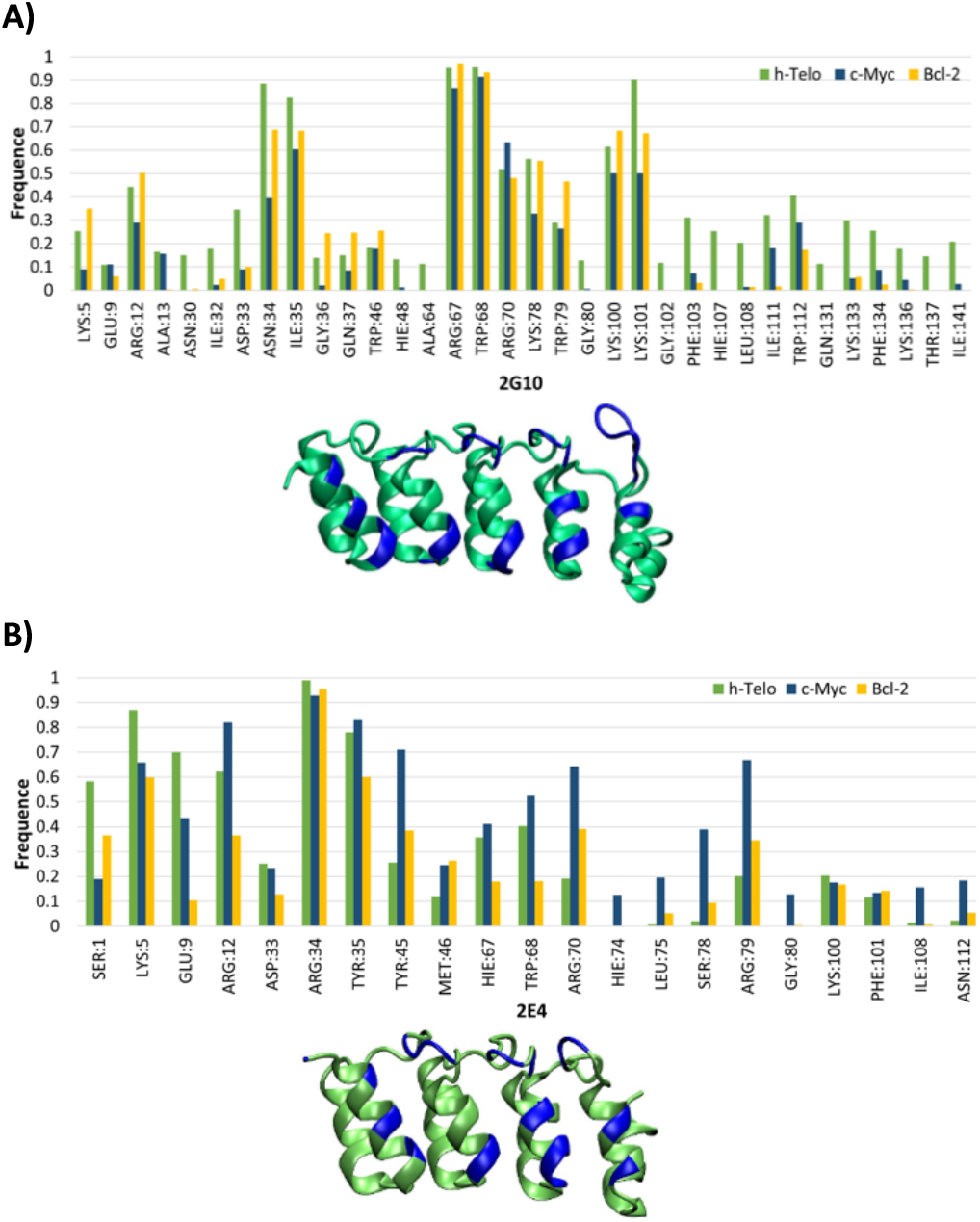
Frequency and location, in blue, of amino acids A) of 2G10 and B) of 2E4 involved in a 3 Å cutoff of G-quadruplex.

Although, the 2G10/G4 complex involves a larger number of residues developing persistent contacts than 2EG/G4, this should not be necessarily correlated to a higher affinity towards G4. Indeed, few residues developing stable and persistent interactions may be regarded as more favorable than an extended weakly interacting region. Furthermore, 2G10 is larger than 2E4, thus the higher number of contacting residues may be also regarded as an obvious statistical effect.

### Alignment between 2E4 and a c-Myc-specific peptide reveals no sequence similarity

Several examples of peptides able to selectively bind G4s are reported in the literature. Usually, they are derived from the DHX36 helicase whose α-helix provides the binding interface with the nucleic acid [39,40]. Notably, Minard et al. [41] designed a specific peptide, DM102 (*PGHLKGRRIGLWYASKQGQKNK)*, which is able to preferentially recognize G4 in the *c-Myc* promoter. Since 2E4 is also specific to *c-Myc*, it is legitimate to ask whether there is a sequence similarity between DM102 and 2E4. It is also important to note that the artificial peptide DM102 has a hydrocarbon staple (*i, i*+7) between residues R8 and S15, which enforces a defined α-helix. Hence, in addition to sequence similarity, one must also look for stable α-helical secondary structure, which is, indeed, a structural motif frequently present in DARPins. Two algorithms were used: Clustal Omega and M-coffee (see Figure S37). Here it is important to specify that Clustal Omega is an individual method of alignment, while M-coffee is an algorithm that combines results from several individual methods [42].

Interestingly, the two algorithms show divergent results. Clustal Omega previews similarity mainly concerning the 2E4 region spanning M46 to V62. The representation of the 3D structure of this region (see Figure S38) reveals a helical arrangement, which could partially support the 2E4 selectivity conditions. However, this region is also common to all DARPins, except for residue 58 (residue 70 following Scholz et al. notation [35]), which is embedded in the similarity region, and residues 45 and 46, which border it. Furthermore, the mutated residues at position 46 and 58 are structurally very distant, suggesting a low quality of the alignment. Conversely, M-Coffee alignment highlights three subunits. Two of them have no significance, the first being located at the previously invalidated region, and the third pertaining to the N-terminal region common to all DARPins. Instead, the second subunit is aligned with the R70-R79 region of DM102, as visually represented in Figure 5 by the transparent shaded area. This observation is also coherent with our MD simulations which indicate an increase of the DARPin/*c-Myc* contact frequency for the residues belonging to this region. Furthermore, from a structural point of view, the R70-R79 region is organized in α-helix motif and is located towards the canonical recognition zone of DARPins. Yet, this region is highly conserved among the DARPins designed by Scholz et al. [35] and only the residues bordering the helix, i.e. R70, S78, and R79, have been mutated. Indeed, when M-Coffee alignment between DM102 and 2G10 the same DARPin region is evidenced (see Figure S39).

**Figure 5.**
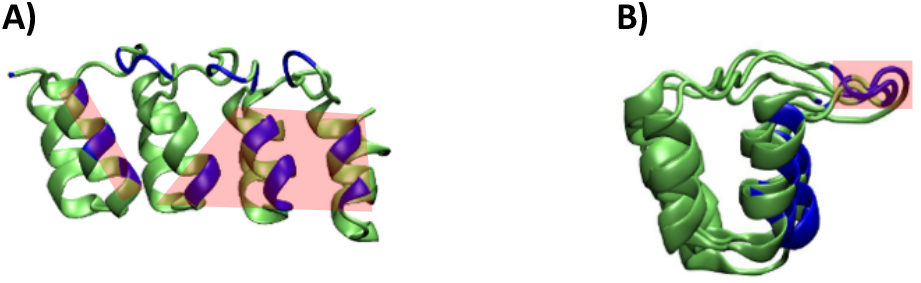
The preferential interaction region, in red, of the 2E4 protein is located at A) the α-helix and B) the loops.

Thus, the search for a conserved sequence between DM102 and 2E4, does not unambiguously justify the selectivity of the DARPin. Going a step further, this could suggest that the recognition of *c-Myc*’s by 2E4 does not necessarily involve sequence similarity between DM102 and 2E4. This is also supported by the fact that the mutations of the wild type sequence as performed by Scholz et al.[35] are mainly concentrated on the peripheral protein loops. Consequently, we decided to focus on structural features which should add up to the rather modest sequence effects and, ultimately, drive the selectivity.

### 2E4 recognizes a particular structural motif of c-Myc

From our MD simulations the local two most important factors should be considered when analyzing the local structural arrangements of the DARPin/G4 the contact region. First, the DARPin canonical interaction zone is not consistently interacting throughout the whole MD simulation. Instead, as highlighted in Figure 4B, other amino acids either located in α-helixes or in peripheral loops develop more persistent interactions.

Furthermore, the analysis of the interaction networks shows that a DARPins/G4 complex is mainly stabilized by electrostatic interaction between positively charged amino acids and the negatively charged backbone of the nucleic acid. In addition, π-cation interactions are also present mainly when the extended conjugated system of a tetrad faces the DARPin. Finally, DARPin associates with G4 through its canonical interaction zone involving the α-helix, but also via interactions mediated by the peripheral loops. The interaction with the loop is most pronounced, but not unique, in the case of *h-Telo*, which in the course of the MD simulation departs from the initial docking pose and slides over the DARPin surface until an interaction between its quartet and the peripheral loops is established at around 150 ns (Figure 6). Interestingly, the electrostatic interactions involving the G4 backbone take place mainly through the G4 external loops rather than the tetrad core. However, this conformation appears as scarcely stable, and as a matter of fact the G4 oscillates and reverts to a more classic interaction mode involving one of its accessible quartets. These observations are also consistent with the frequency distribution reported in Figure 3 and explain the specific behavior of *h-Telo*, which due to its high mobility spans different interaction poses and develops rather non-specific contacts with a high number of 2E4 and 2G10 residues.

**Figure 6.**
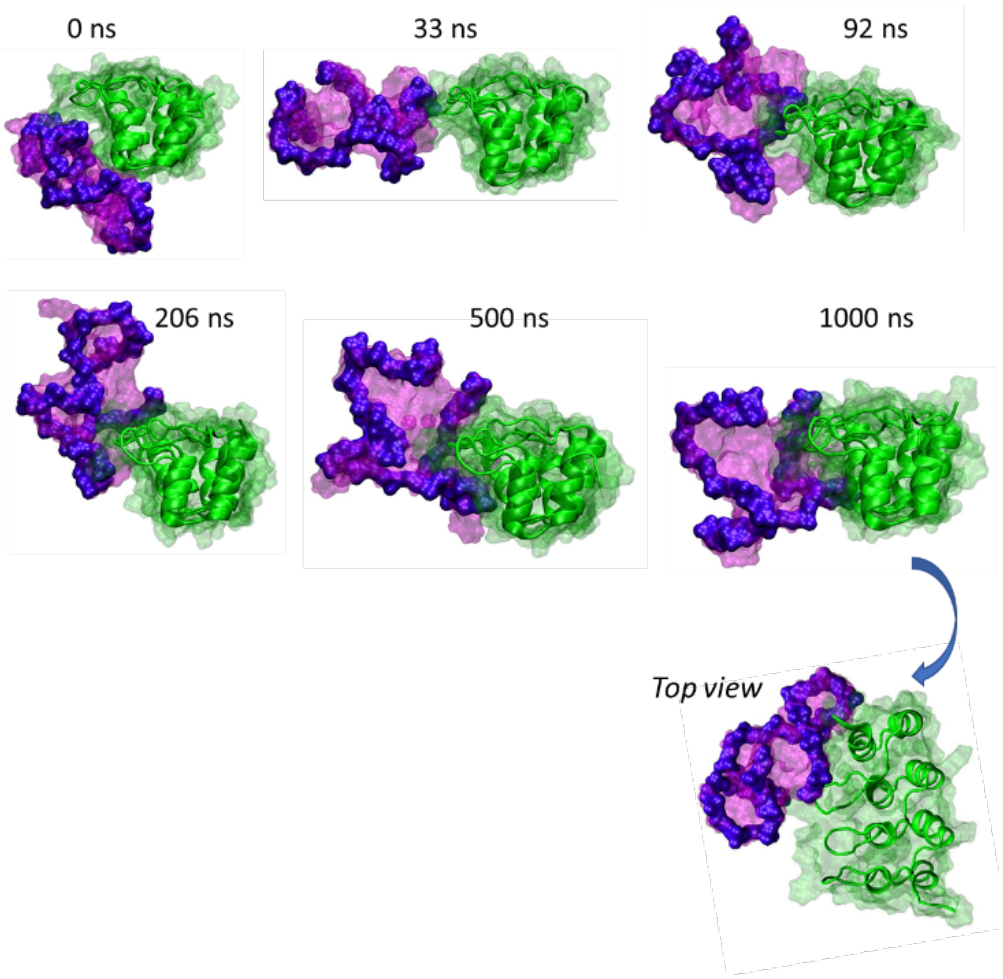
Screenshots of the dynamics of a 2E4/*h-Telo* complex. The *h-Telo* quartet is initially oriented towards the a-helix. But within the first 150 ns, the G-quadruplex rotates and then reorients to interact with the protein loops. During the dynamics, the movements of the G-quadruplex do not affect the protein/G-quartet loops interaction.

Thus, the interaction mode involving a quartet is not leading to a specific recognition mode. Hence, interactions between *c-Myc* or *Bcl-2* and DARPin which would be driven by the G4s quartets (Figure 7) will most probably be trapped in an non-specific recognition and cannot be used to infer on the specific recognition. On the contrary, specificity may be established when DARPins interact mainly with the nucleic acid backbone. The behavior of *Bcl-2*’s, which interacts in a similar non-specific way with 2E4, confirms nicely this statement. Indeed, despite different initial conditions, the G4 again positions itself exposing a quartet to the 2E4 DARPin interaction region. On the contrary, the interaction with 2G10 leads to the exposure of the nucleic acid backbone to the contact region of DARPin and hence, to a selective recognition. As a matter of fact, these results are also coherent with the contact frequency analysis showing that *Bcl-2* interaction with 2G10 is mostly driven by highly conserved and persistent aminoacids.

**Figure 7.**
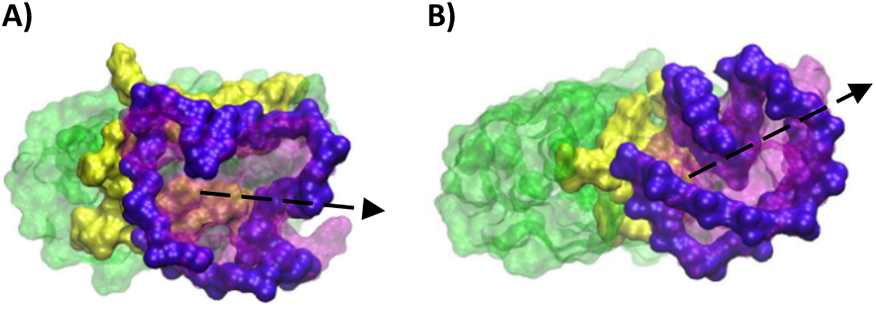
Non-specific ‘nucleotide face’ interaction found in cluster of the A) 2E4*/h-Telo* pose 1-1 (77.23% of the MDs) and B) 2E4/*Bcl-2* pose 1-1 (79.01% of the MDs) complexes.

### 2E4 recognizes a peculiar structural motif of c-Myc

*c-Myc* is the G4 more consistently promoting an interaction via its backbone (Figure 8). This, in turn, could also point to a greater specificity of its recognition, although *c-Myc*’s is able to interact via its backbone with both 2E4 and 2G10. Thus, to further justify the selectivity of 2E4 a structural motif specifically recognized by this DARPin should be identified.

**Figure 8.**
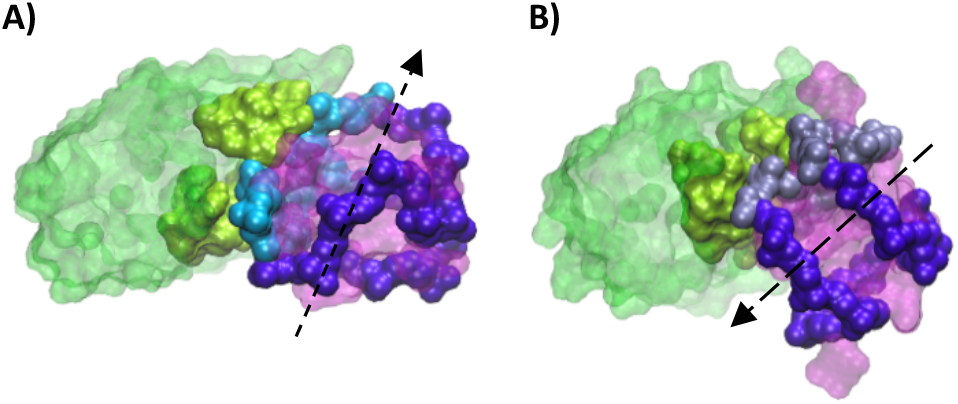
*c-Myc* G-quadruplex interacts preferentially through its backbone A) in the dynamics resulting from the most stable docking pose and B) in the dynamics resulting from a less favorable pose in which the G-quadruplex reorients itself to interact in a pose like the most stable pose found by docking. The previously identified high frequency nucleotides are colored A) in cyan and B) in steel blue, respectively.

The main factor that could lead to a recognized structural motif includes the presence of a backbone folding involving the nucleotides most frequently in contact with the DARPin. This feature can be easily assessed by clustering the MD simulation while checking the maintenance of the interaction patterns in the most populated clusters. By highlighting the E24 highest frequency contact nucleotides, i.e. G6 to A12, A21, and A22, we see that they are involved in the interaction with the protein for the two most important clusters (Figure 8 and 9A, 9B). However, the two clusters differ by a rotation of about 180° of the G4 on the protein surface (pose 1-1: 78.47% of the MDs and pose 6-4: 79.55% of the MDs), as shown in Figure 10. Nonetheless, a well conversed structural motif is evidenced, determined by the folding of the G4 backbone into a U-shaped loop with an extruded nucleotide, further completed by a horizontal extension to the right, and overlaid by a dangling segment (Figure 9). Interestingly, all the structural characteristics are well evidenced in the most populated cluster, while in the secondary structures their identification remains more elusive. Indeed, if the linear extension remains evident, as well as the extruded nucleotide, the U-shaped loop and the appendix are more scarcely visible.

**Figure 9.**
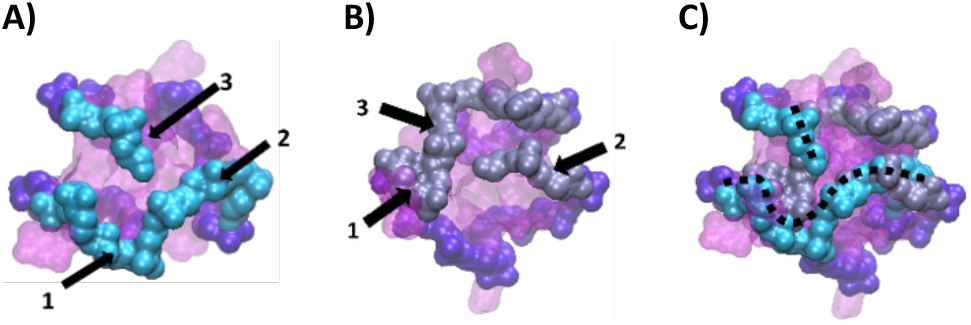
Face view of the *c-Myc* G-quadruplex interacting with 2E4 A) in the dynamics resulting from the most stable docking pose (pose 1-1) and B) in the dynamics resulting from a less favorable pose (pose 6-4): a rotation of 180 degrees with respect to A). Three elements can be identified on each: 1. a U-shaped-loop with an extruded nucleotide, 2. a linear extension and 3. an appendage. The superposition of the two interaction poses C), identify the recognized motif in dotted line.

The amino acids located in the interaction site are also conserved between the two most populated clusters (Figure 10). Residues Y45, R70, and R79 organizes around the loop at an average distance of 3 Å, while Y35 is oriented towards the appendix and R34 points towards the linear extension. At a slightly higher distance of around 5 Å, M46 is interacting with the U-shaped loop, W68 is oriented towards the appendix, while R12 and D33 flank the linear extension. In addition, S78 stays close to the extruded nucleotide, probably assuring a further stabilization.

**Figure 10.**
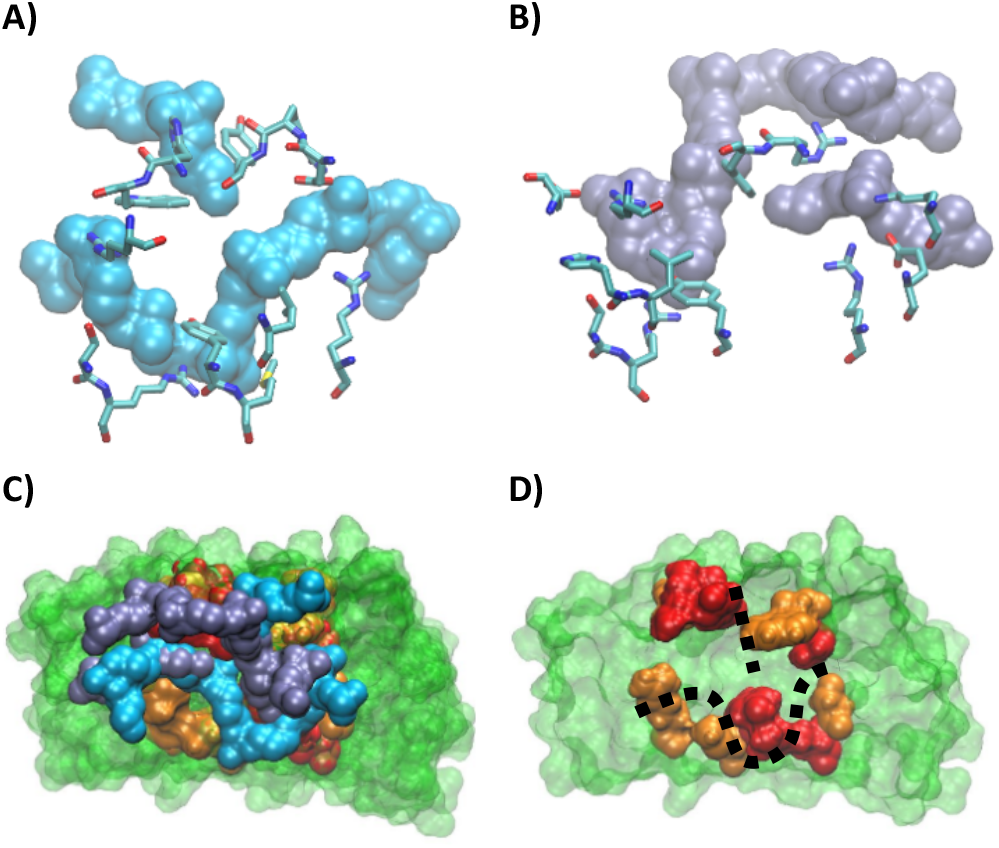
Protein residue involved in the interaction with the c-Myc structural motif A) in the dynamics resulting from the most stable docking pose (pose 1-1) and B) in the dynamics resulting from a less favorable pose (pose 6-4). C) Superposition of the complexes of the two clusters. D) Interaction site of 2E4 with c-Myc; residues at 3 Å are in red and those at 5 Å are in orange.

**Figure 13.**
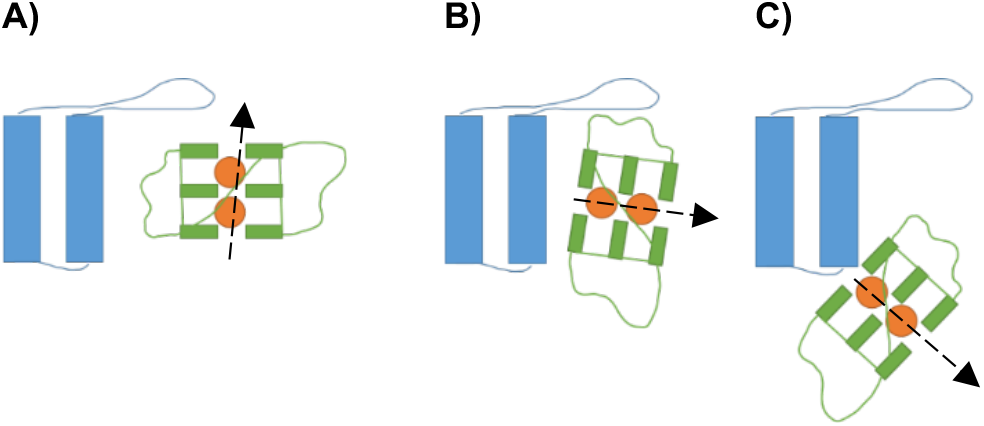
Schematic representation of the three types of poses selected from the docking. There is interaction type A) backbone, B) nucleotide face and C) protein bottom zone.

By superposing the two most populated clusters of the 2E4/*c-Myc* complexes a similar positioning of the G4 on the protein site is also observed, which is again consistent with the recognized backbone-based structural motif (Figure 10D). Finally, the increase of the contact frequency observed in the analysis of the MD simulations correlates well with the specific recognition of *c-Myc* by 2E4. Indeed, the region presenting the highest increase in the contact frequency corresponds to the residues recognizing the U-shaped loop and the extruded nucleotide. This, together with the similarity of the interaction pattern found in the superposition of the two G4s poses, further validates the hypothesis that the selective recognition of *c-Myc* by 2E4 is driven by the structural motif we have identified. Because of this structural-based recognition, and the often-invoked analogy between DARPin and antibodies, it is tempting to characterize this interaction pattern as an epitope/paratope recognition. Here the paratope-like element being the 2E4 interaction site, and the epitope-like region the G4 structural motif identified for *c-Myc*.

## Conclusions

Our results highlight two modes of interaction for DARPins/G4 complexes. The first one is a non-specific recognition that is established when G4 interacts through its guanine tetrad, or through peripheral nucleotides π-stacked with the tetrads. The second binding mode is driven by the specific recognition of the conformation of the G4 backbone and leads to a DARPin/G4 paratope/epitope like recognition. This specific mode, which we have identified for the 2E4/*C-myc* complex is based on a peculiar folding motif of the G4 backbone and presents a U-shaped loop with a linear extension and an overhanging short appendix. Consequently, a large extension of the U-shaped loop, also including extruded nucleotides should enhance selective recognition of the G4s. Conversely, the identification of backbone-based recognition motifs could also improve the rational design of DARPins. Indeed, the quest for selective G4 ligands has a tremendous significance, especially in the proposition of specific anticancer or antiviral agents. Our results, and the first identification of paratope/epitope specific structural recognition may lead to significant development in the design of potentially therapeutics peptides targeting specific G4 arrangements.

## Experimental Section

### Structure of G4 and reconstruction of DARPins

The structure of the *h-Telo* G4 was retrieved from PDB data bank 1KF1 [43], as well as that of the *c-Myc* (1XAV) [44] and *Bcl-2 (*6ZX7) [45]. The sequences of 2E4 and 2G10 DARPins were obtained from the Supplementary Information of [35] and their structure reconstructed with the SWISS-MODEL server [46]. 2E4 were reconstructed based on high similarity with the PDB entry 2CH4 [47] and 2G10 was reconstructed on the basis of the 1SVX structure [48] similarity.

### Sequence alignment

DM102 peptide and DARPins sequences were aligned using the Clustal Omega on EBI server [49] and the M-coffee server [50], using their default parameters.

### Docking and selection of initial structures

The reconstructed DARPins and G4 were loaded onto the HADDOCK server [51] to perform protein/nucleic acid docking while searching the entire protein and the entire G4 structure and using the standard HADDOCK parameters. Three poses were selected from the docking results, always including the most favorable one. The selection was based on the relative position of G4 with respect to the DARPin. The three poses correspond to an interaction with the G4 backbone, an interaction with the tetrads facing the nucleotides and an interaction developed in a peripheral region of the DARPin. This choice allowed to assure a significant sampling of a complex conformational space, also including rather unfavorable interaction areas, such as the one corresponding to the peripheral binding.

### Molecular dynamics simulations

MD simulations has been performed for 2E4 and 2G10 interacting with *c-Myc, Bcl-2* and *h-Telo*. Three poses for each complex have been used as starting conditions. Each system was calculated in two independent replicates of 1 µs each, thus a total of 36 simulations of DARPin/G4complexes have been performed. In addition. simulations of the free G4s and DARPin have also been obtained as a control. All simulations have been run using the NAMD software [52] with the Amber parm99 force field [53] including the bsc1 correction [54] for nucleic acids. An octahedral box of TIP3P [55] water was used to solvate the systems, using periodic boundary condition (PBC). All the calculations were performed in the isothermal and isobaric (NPT) ensemble at a temperature of 300K and a pressure of 1 atm. A minimal concentration of K+ ions was added to assure charge equilibration. Hydrogen Mass Repartitioning (HMR) [56] was consistently used, allowing, in combination with the Rattle and Shake algorithms [57], a timestep of 4 fs to integrate Newton’s equations of motion. Finally, the trajectories were analyzed and visualized with VMD [58], as well as a dedicated script to retrieve G4 structural parameters [59], while CPPTRAJ [60] was used for clustering.

## Acknowledgements

All the calculations were performed on the local cluster of the LPCT and on the regional cluster ExpLor of the University of Lorraine. CNRS and Université de Lorraine and Université Paris Cité, France, and Università di Palermo and from the Ministero dell’Università e Ricerca Scientifica e Tecnologica, are thanked for their support. T.M. thanks University of Palermo for funding a joint Ph.D. program. A.M. thanks ANR (Agence Nationale de la Recherche) and CGI (Commissariat à l’Investissement d’Avenir) for their financial support of this work through Labex SEAM (Science and Engineering for Advanced Materials and devices) ANR 11 LABX 086 and ANR 11 IDEX 05 02. The support of the IdEx “Université Paris 2019” ANR-18-IDEX-0001 and of the Platform P3MB is gratefully acknowledged.

## Entry for the Table of Contents

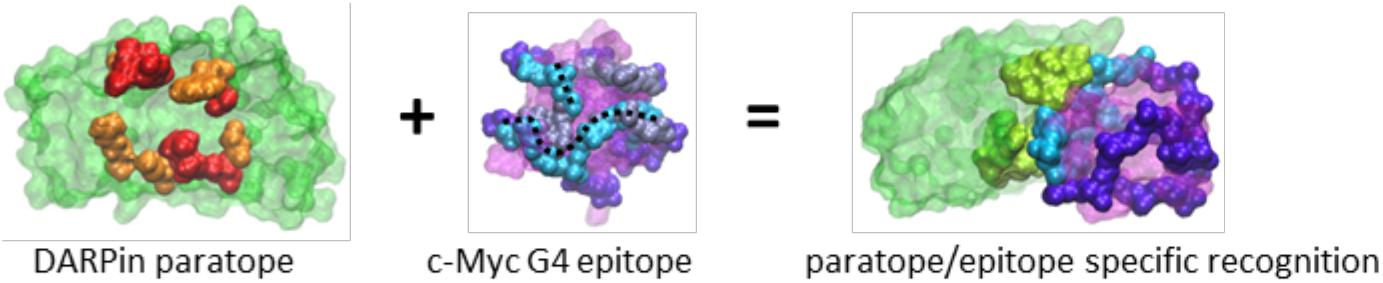

DARPins are proteins able to specifically recognize G-quadruplexes. Their small size combined with their easy design makes them a good competitor to antibodies for the identification and localization of G-quadruplex. Using molecular dynamics calculations, we show that the selectivity of DARPins towards G-quadruplexes is achieved by the recognition of a structural motif adopted by the DNA filament folding.

Institute and/or researcher Twitter usernames: ((optional))

## References

1. Lipps, H.J., and Rhodes, D. (2009) G-quadruplex structures: in vivo evidence and function. Trends in Cell Biology, 19 (8), 414–422.

2. Bochman, M.L., Paeschke, K., and Zakian, V.A. (2012) DNA secondary structures: stability and function of G-quadruplex structures. Nature Reviews Genetics, 13 (11), 770–780.

3. Kwok, C.K., and Merrick, C.J. (2017) G-Quadruplexes: Prediction, Characterization, and Biological Application. Trends in Biotechnology, 35 (10), 997–1013.

4. Spiegel, J., Adhikari, S., and Balasubramanian, S. (2020) The Structure and Function of DNA G-Quadruplexes. Trends in Chemistry, 2 (2), 123–136.

5. David, A.P., Margarit, E., Domizi, P., Banchio, C., Armas, P., and Calcaterra, N.B. (2016) G-quadruplexes as novel cis-elements controlling transcription during embryonic development. Nucleic Acids Research, 44 (9), 4163–4173.

6. Hänsel-Hertsch, R., Beraldi, D., Lensing, S. v, Marsico, G., Zyner, K., Parry, A., di Antonio, M., Pike, J., Kimura, H., Narita, M., Tannahill, D., and Balasubramanian, S. (2016) G-quadruplex structures mark human regulatory chromatin. Nature Genetics, 48 (10), 1267–1272.

7. Moruno-Manchon, J.F., Lejault, P., Wang, Y., McCauley, B., Honarpisheh, P., Morales Scheihing, D.A., Singh, S., Dang, W., Kim, N., Urayama, A., Zhu, L., Monchaud, D., McCullough, L.D., and Tsvetkov, A.S. (2020) Small-molecule G-quadruplex stabilizers reveal a novel pathway of autophagy regulation in neurons. Elife, 9.

8. Simone, R., Fratta, P., Neidle, S., Parkinson, G.N., and Isaacs, A.M. (2015) G-quadruplexes: Emerging roles in neurodegenerative diseases and the non-coding transcriptome. FEBS Letters, 589 (14), 1653–1668.

9. Biffi, G., Tannahill, D., Miller, J., Howat, W.J., and Balasubramanian, S. (2014) Elevated Levels of G-Quadruplex Formation in Human Stomach and Liver Cancer Tissues. PLoS ONE, 9 (7), e102711.

10. Carvalho, J., Mergny, J.-L., Salgado, G.F., Queiroz, J.A., and Cruz, C. (2020) G-quadruplex, Friend or Foe: The Role of the G-quartet in Anticancer Strategies. Trends in Molecular Medicine, 26 (9), 848–861.

11. Xu, Y.-Z., Jenjaroenpun, P., Wongsurawat, T., Byrum, S.D., Shponka, V., Tannahill, D., Chavez, E.A., Hung, S.S., Steidl, C., Balasubramanian, S., Rimsza, L.M., and Kendrick, S. (2020) Activation-induced cytidine deaminase localizes to G-quadruplex motifs at mutation hotspots in lymphoma. NAR Cancer, 2 (4).

12. Kosiol, N., Juranek, S., Brossart, P., Heine, A., and Paeschke, K. (2021) G-quadruplexes: a promising target for cancer therapy. Molecular Cancer, 20 (1), 40.

13. Kumari, N., Vartak, S. v., Dahal, S., Kumari, S., Desai, S.S., Gopalakrishnan, V., Choudhary, B., and Raghavan, S.C. (2019) G-quadruplex Structures Contribute to Differential Radiosensitivity of the Human Genome. iScience, 21, 288–307.

14. Bielskutė, S., Plavec, J., and Podbevšek, P. (2019) Impact of Oxidative Lesions on the Human Telomeric G-Quadruplex. J Am Chem Soc, 141 (6), 2594–2603.

15. Fleming, A.M., Zhu, J., Ding, Y., and Burrows, C.J. (2017) 8-Oxo-7,8-dihydroguanine in the Context of a Gene Promoter G-Quadruplex Is an On–Off Switch for Transcription. ACS Chemical Biology, 12 (9), 2417–2426.

16. Kosiol, N., Juranek, S., Brossart, P., Heine, A., and Paeschke, K. (2021) G-quadruplexes: a promising target for cancer therapy. Molecular Cancer, 20 (1), 40.

17. Hognon, C., Miclot, T., García-Iriepa, C., Francés-Monerris, A., Grandemange, S., Terenzi, A., Marazzi, M., Barone, G., and Monari, A. (2020) Role of RNA Guanine Quadruplexes in Favoring the Dimerization of SARS Unique Domain in Coronaviruses. The Journal of Physical Chemistry Letters, 11 (14), 5661–5667.

18. Francés-Monerris, A., Hognon, C., Miclot, T., GarcíaIriepa, C., Iriepa, I., Terenzi, A., Grandemange, S., Barone, G., Marazzi, M., and Monari, A. (2020) Molecular Basis of SARS-CoV-2 Infection and Rational Design of Potential Antiviral Agents: Modeling and Simulation Approaches. Journal of Proteome Research, 19 (11), 4291–4315.

19. Badmalia, M., Meier-Stephenson, V., Balderas Figueroa, G., D’Souza, S., Coffin, C., and Patel, T.R. (2022) G-Quadruplex in hepatitis B virus. Biophysical Journal, 121 (3), 66a.

20. Zhao, C., Qin, G., Niu, J., Wang, Z., Wang, C., Ren, J., and Qu, X. (2021) Targeting RNA G-Quadruplex in SARS-CoV-2: A Promising Therapeutic Target for COVID-19? Angewandte Chemie International Edition, 60 (1), 432–438.

21. Wang, S.-R., Zhang, Q.-Y., Wang, J.-Q., Ge, X.-Y., Song, Y.-Y., Wang, Y.-F., Li, X.-D., Fu, B.-S., Xu, G.-H., Shu, B., Gong, P., Zhang, B., Tian, T., and Zhou, X. (2016) Chemical Targeting of a G-Quadruplex RNA in the Ebola Virus L Gene. Cell Chemical Biology, 23 (9), 1113–1122.

22. Saranathan, N., and Vivekanandan, P. (2019) G-Quadruplexes: More Than Just a Kink in Microbial Genomes. Trends in Microbiology, 27 (2), 148–163.

23. Chen, M.C., Tippana, R., Demeshkina, N.A., Murat, P., Balasubramanian, S., Myong, S., and Ferré-D’Amaré, A.R. (2018) Structural basis of G-quadruplex unfolding by the DEAH/RHA helicase DHX36. Nature, 558 (7710), 465–469.

24. Pietras, Z., Wojcik, M.A., Borowski, L.S., Szewczyk, M., Kulinski, T.M., Cysewski, D., Stepien, P.P., Dziembowski, A., and Szczesny, R.J. (2018) Dedicated surveillance mechanism controls G-quadruplex forming non-coding RNAs in human mitochondria. Nature Communications, 9 (1), 2558.

25. Darnell, J.C., Jensen, K.B., Jin, P., Brown, V., Warren, S.T., and Darnell, R.B. (2001) Fragile X Mental Retardation Protein Targets G Quartet mRNAs Important for Neuronal Function. Cell, 107 (4), 489–499.

26. Farine, G., Migliore, C., Terenzi, A., lo Celso, F., Santoro, A., Bruno, G., Bonsignore, R., and Barone, G. (2021) On the G-Quadruplex Binding of a New Class of Nickel(II), Copper(II), and Zinc(II) Salphen-Like Complexes. European Journal of Inorganic Chemistry, 2021 (14), 1332–1336.

27. Dhamodharan, V., and Pradeepkumar, P.I. (2019) Specific Recognition of Promoter G-Quadruplex DNAs by Small Molecule Ligands and Light-up Probes. ACS Chemical Biology, acschembio.9b00475.

28. Drygin, D., Siddiqui-Jain, A., O’Brien, S., Schwaebe, M., Lin, A., Bliesath, J., Ho, C.B., Proffitt, C., Trent, K., Whitten, J.P., Lim, J.K.C., von Hoff, D., Anderes, K., and Rice, W.G. (2009) Anticancer Activity of CX-3543: A Direct Inhibitor of rRNA Biogenesis. Cancer Research, 69 (19), 7653–7661.

29. Dutta, D., Debnath, M., Müller, D., Paul, R., Das, T., Bessi, I., Schwalbe, H., and Dash, J. (2018) Cell penetrating thiazole peptides inhibit c-MYC expression via site-specific targeting of c-MYC G-quadruplex. Nucleic Acids Research, 46 (11), 5355–5365.

30. Javadekar, S.M., Nilavar, N.M., Paranjape, A., Das, K., and Raghavan, S.C. (2020) Characterization of G-quadruplex antibody reveals differential specificity for G4 DNA forms. DNA Research, 27 (5).

31. Henderson, A., Wu, Y., Huang, Y.C., Chavez, E.A., Platt, J., Johnson, F.B., Brosh, R.M., Sen, D., and Lansdorp, P.M. (2017) Detection of G-quadruplex DNA in mammalian cells. Nucleic Acids Research, 45 (10), 6252–6252.

32. Javadekar, S.M., Nilavar, N.M., Paranjape, A., Das, K., and Raghavan, S.C. (2020) Characterization of G-quadruplex antibody reveals differential specificity for G4 DNA forms. DNA Research, 27 (5).

33. Liu, H.-Y., Zhao, Q., Zhang, T.-P., Wu, Y., Xiong, Y.-X., Wang, S.-K., Ge, Y.-L., He, J.-H., Lv, P., Ou, T.-M., Tan, J.-H., Li, D., Gu, L.-Q., Ren, J., Zhao, Y., and Huang, Z.-S. (2016) Conformation Selective Antibody Enables Genome Profiling and Leads to Discovery of Parallel G-Quadruplex in Human Telomeres. Cell Chemical Biology, 23 (10), 1261–1270.

34. Fernando, H., Rodriguez, R., and Balasubramanian, S. (2008) Selective Recognition of a DNA G-Quadruplex by an Engineered Antibody. Biochemistry, 47 (36), 9365–9371.

35. Scholz, O., Hansen, S., and Plückthun, A. (2014) G-quadruplexes are specifically recognized and distinguished by selected designed ankyrin repeat proteins. Nucleic Acids Research, 42 (14), 9182–9194.

36. Mittl, P.R., Ernst, P., and Plückthun, A. (2020) Chaperone-assisted structure elucidation with DARPins. Current Opinion in Structural Biology, 60, 93–100.

37. Plückthun, A. (2015) Designed Ankyrin Repeat Proteins (DARPins): Binding Proteins for Research, Diagnostics, and Therapy. Annual Review of Pharmacology and Toxicology, 55 (1), 489–511.

38. Stumpp, M.T., Binz, H.K., and Amstutz, P. (2008) DARPins: A new generation of protein therapeutics. Drug Discovery Today, 13 (15–16), 695–701.

39. Heddi, B., Cheong, V.V., Martadinata, H., and Phan, A.T. (2015) Insights into G-quadruplex specific recognition by the DEAH-box helicase RHAU: Solution structure of a peptide–quadruplex complex. Proceedings of the National Academy of Sciences, 112 (31), 9608–9613.

40. Yaneva, M.Y., Cheong, V.V., Cheng, J.K., Lim, K.W., and Phan, A.T. (2020) Stapling a G-quadruplex specific peptide. Biochemical and Biophysical Research Communications, 531 (1), 62–66.

41. Minard, A., Morgan, D., Raguseo, F., di Porzio, A., Liano, D., Jamieson, A.G., and di Antonio, M. (2020) A short peptide that preferentially binds c-MYC G-quadruplex DNA. Chemical Communications, 56 (63), 8940–8943.

42. Wallace, I.M. (2006) M-Coffee: combining multiple sequence alignment methods with T-Coffee. Nucleic Acids Research, 34 (6), 1692–1699.

43. Parkinson, G.N., Lee, M.P.H., and Neidle, S. (2002) Crystal structure of parallel quadruplexes from human telomeric DNA. Nature, 417 (6891).

44. Ambrus, A., Chen, D., Dai, J., Jones, R.A., and Yang, D. (2005) Solution Structure of the Biologically Relevant G-Quadruplex Element in the Human c-MYC Promoter. Implications for G-Quadruplex Stabilization. Biochemistry, 44 (6), 2048–2058.

45. Bielskutė, S., Plavec, J., and Podbevšek, P. (2021) Oxidative lesions modulate G-quadruplex stability and structure in the human BCL2 promoter. Nucleic Acids Research, 49 (4), 2346–2356.

46. Waterhouse, A., Bertoni, M., Bienert, S., Studer, G., Tauriello, G., Gumienny, R., Heer, F.T., de Beer, T.A.P., Rempfer, C., Bordoli, L., Lepore, R., and Schwede, T. (2018) SWISS-MODEL: homology modelling of protein structures and complexes. Nucleic Acids Research, 46 (W1), W296–W303.

47. Grubisha, O., Kaminska, M., Duquerroy, S., Fontan, E., Cordier, F., Haouz, A., Raynal, B., Chiaravalli, J., Delepierre, M., Israël, A., Véron, M., and Agou, F. (2010) DARPin-Assisted Crystallography of the CC2-LZ Domain of NEMO Reveals a Coupling between Dimerization and Ubiquitin Binding. Journal of Molecular Biology, 395 (1), 89–104.

48. Binz, H.K., Amstutz, P., Kohl, A., Stumpp, M.T., Briand, C., Forrer, P., Grütter, M.G., and Plückthun, A. (2004) High-affinity binders selected from designed ankyrin repeat protein libraries. Nature Biotechnology, 22 (5), 575–582.

49. Madeira, F., Park, Y. mi, Lee, J., Buso, N., Gur, T., Madhusoodanan, N., Basutkar, P., Tivey, A.R.N., Potter, S.C., Finn, R.D., and Lopez, R. (2019) The EMBL-EBI search and sequence analysis tools APIs in 2019. Nucleic Acids Research, 47 (W1), W636–W641.

50. Moretti, S., Armougom, F., Wallace, I.M., Higgins, D.G., Jongeneel, C. v., and Notredame, C. (2007) The M-Coffee web server: a meta-method for computing multiple sequence alignments by combining alternative alignment methods. Nucleic Acids Research, 35 (Web Server), W645–W648.

51. van Zundert, G.C.P., Rodrigues, J.P.G.L.M., Trellet, M., Schmitz, C., Kastritis, P.L., Karaca, E., Melquiond, A.S.J., van Dijk, M., de Vries, S.J., and Bonvin, A.M.J.J. (2016) The HADDOCK2.2 Web Server: User-Friendly Integrative Modeling of Biomolecular Complexes. Journal of Molecular Biology, 428 (4), 720–725.

52. Phillips, J.C., Braun, R., Wang, W., Gumbart, J., Tajkhorshid, E., Villa, E., Chipot, C., Skeel, R.D., Kalé, L., and Schulten, K. (2005) Scalable molecular dynamics with NAMD. Journal of Computational Chemistry, 26 (16).

53. Cornell, W.D., Cieplak, P., Bayly, C.I., Gould, I.R., Merz, K.M., Ferguson, D.M., Spellmeyer, D.C., Fox, T., Caldwell, J.W., and Kollman, P.A. (1995) A Second Generation Force Field for the Simulation of Proteins, Nucleic Acids, and Organic Molecules. J Am Chem Soc, 117 (19), 5179–5197.

54. Ivani, I., Dans, P.D., Noy, A., Pérez, A., Faustino, I., Hospital, A., Walther, J., Andrio, P., Goñi, R., Balaceanu, A., Portella, G., Battistini, F., Gelpí, J.L., González, C., Vendruscolo, M., Laughton, C.A., Harris, S.A., Case, D.A., and Orozco, M. (2016) Parmbsc1: a refined force field for DNA simulations. Nature Methods, 13 (1), 55–58.

55. Mark, P., and Nilsson, L. (2001) Structure and Dynamics of the TIP3P, SPC, and SPC/E Water Models at 298 K. The Journal of Physical Chemistry A, 105 (43).

56. Hopkins, C.W., le Grand, S., Walker, R.C., and Roitberg, A.E. (2015) Long-Time-Step Molecular Dynamics through Hydrogen Mass Repartitioning. Journal of Chemical Theory and Computation, 11 (4).

57. Miyamoto, S., and Kollman, P.A. (1992) Settle: An analytical version of the SHAKE and RATTLE algorithm for rigid water models. Journal of Computational Chemistry, 13 (8).

58. Humphrey, W., Dalke, A., and Schulten, K. (1996) VMD: Visual molecular dynamics. Journal of Molecular Graphics, 14 (1).

59. Tsvetkov, V., Pozmogova, G., and Varizhuk, A. (2016) The systematic approach to describing conformational rearrangements in G-quadruplexes. Journal of Biomolecular Structure and Dynamics, 34 (4).

60. Roe, D.R., and Cheatham, T.E. (2013) PTRAJ and CPPTRAJ: Software for Processing and Analysis of Molecular Dynamics Trajectory Data. Journal of Chemical Theory and Computation, 9 (7), 3084–3095.

