## Supplementary Information for "G-quadruplex recognition by DARPIns through epitope/paratope analogy"

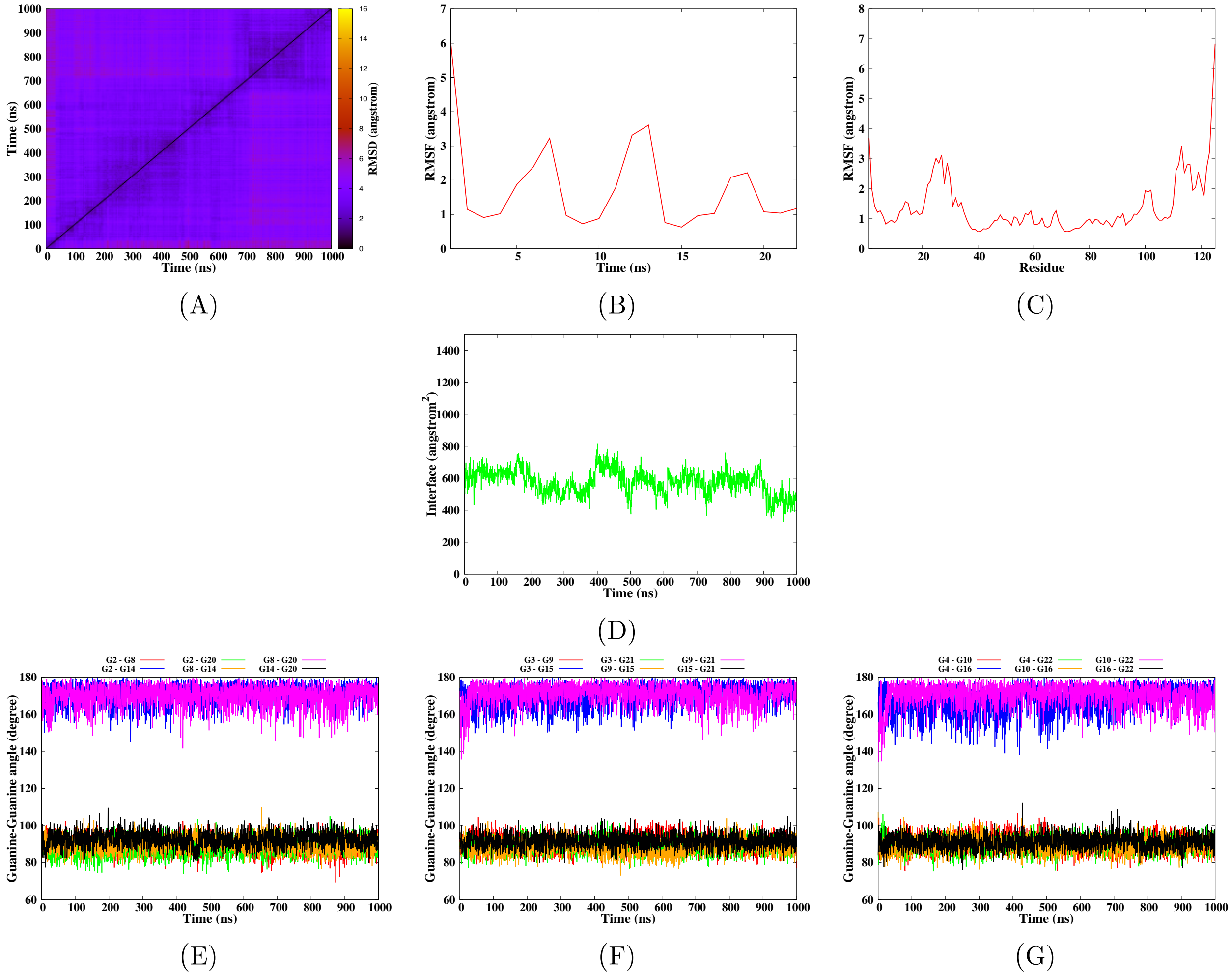


**Figure S1** – Simulation of the h-Telo G-quadruplex DNA in interaction with 2E4 according to the model 1-1, run 1. The convergence of the simulation is given by the RMSD-2D map of the DNA-Protein complex (A). The mobility of the DNA and protein residues is given by their root mean square fluctuation (B-C). Surface of the interaction interface between the protein and G-quadruplex (D). Finally the structural parameters of the G-quadruplex are given by the angles between the guanines for each tetrad (E-G).

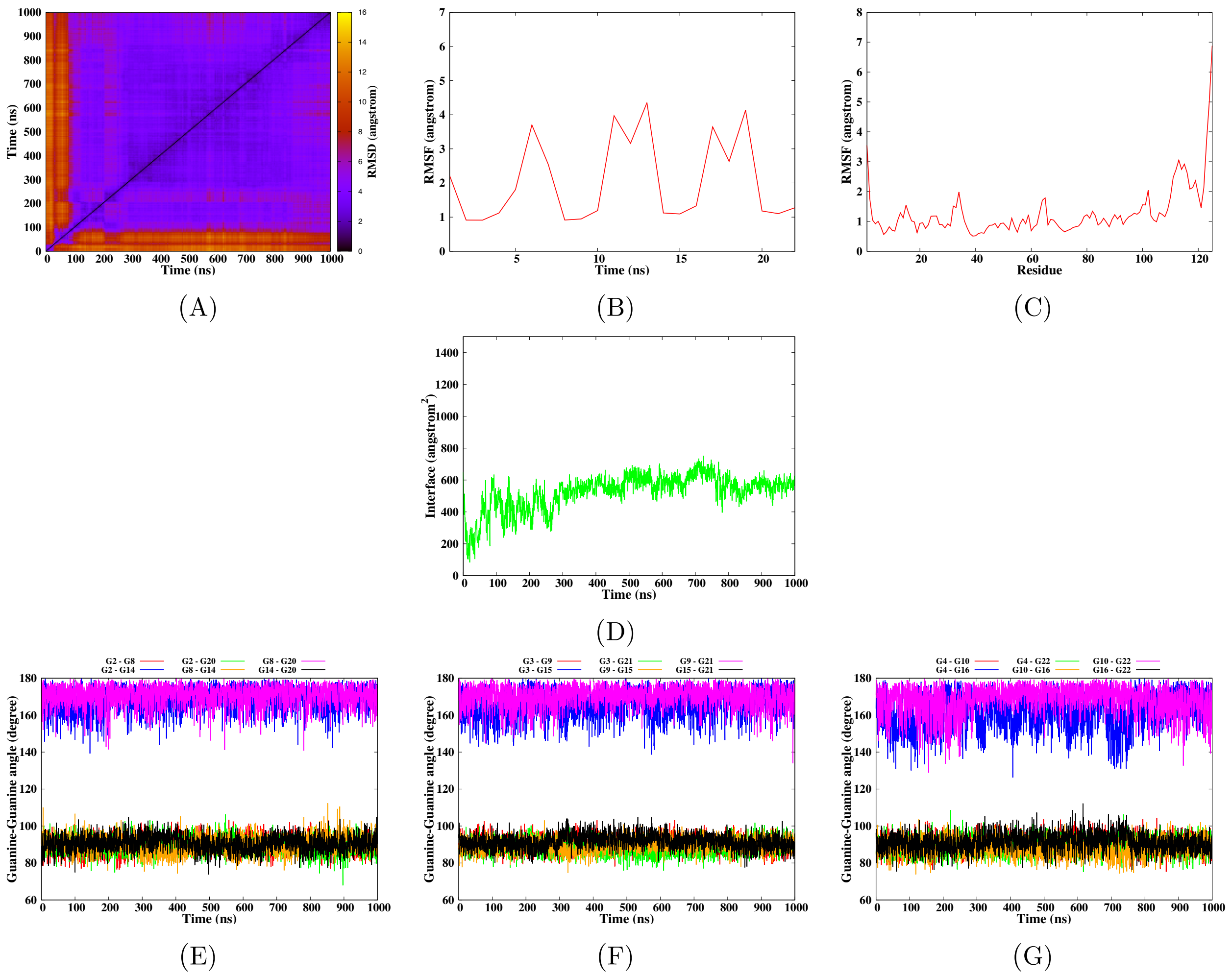


**Figure S2** – Simulation of the h-Telo G-quadruplex DNA in interaction with 2E4 according to the model 1-1, run 2. The convergence of the simulation is given by the RMSD-2D map of the DNA-Protein complex (A). The mobility of the DNA and protein residues is given by their root mean square fluctuation (B-C). Surface of the interaction interface between the protein and G-quadruplex (D). Finally the structural parameters of the G-quadruplex are given by the angles between the guanines for each tetrad (E-G).

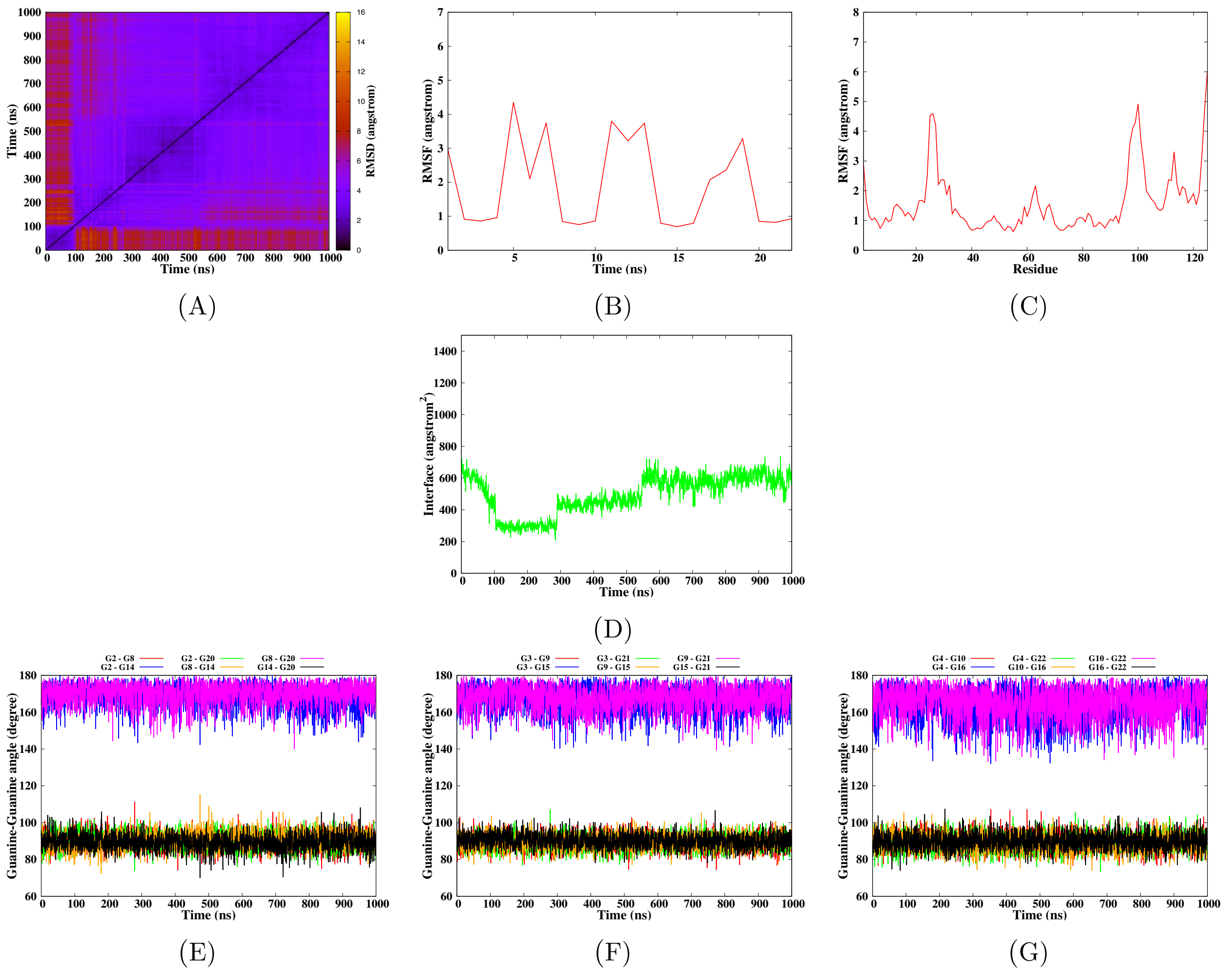


**Figure S3** – Simulation of the h-Telo G-quadruplex DNA in interaction with 2E4 according to the model 6-1, run 1. The convergence of the simulation is given by the RMSD-2D map of the DNA-Protein complex (A). The mobility of the DNA and protein residues is given by their root mean square fluctuation (B-C). Surface of the interaction interface between the protein and G-quadruplex (D). Finally the structural parameters of the G-quadruplex are given by the angles between the guanines for each tetrad (E-G).

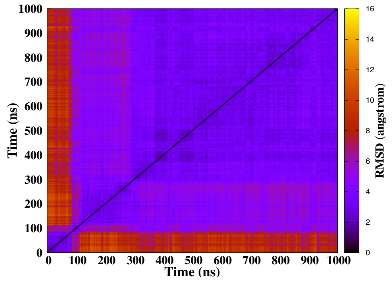


(A)

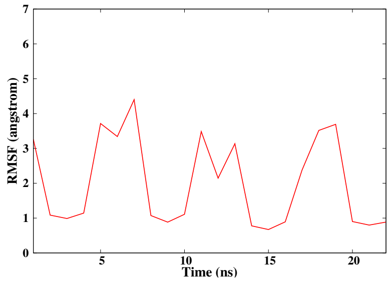


(B)

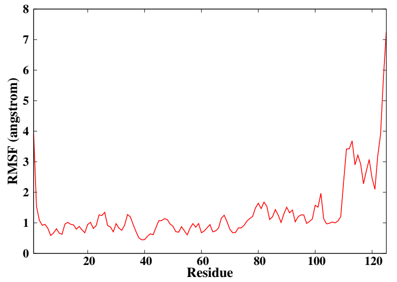


(C)

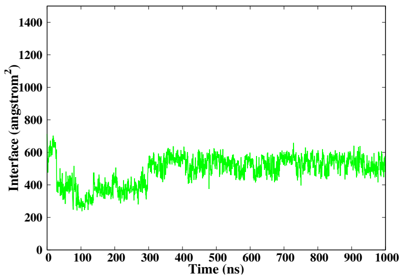


(D)

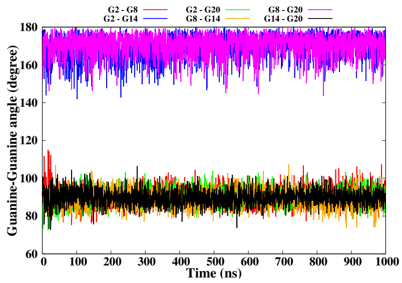


(E)

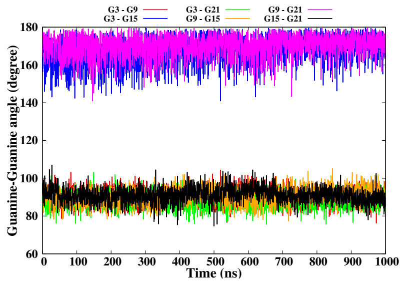


(F)

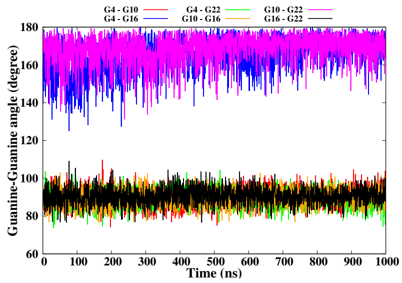


(G)

**Figure S4** – Simulation of the h-Telo G-quadruplex DNA in interaction with 2E4 according to the model 6-1, run 2. The convergence of the simulation is given by the RMSD-2D map of the DNA-Protein complex (A). The mobility of the DNA and protein residues is given by their root mean square fluctuation (B-C). Surface of the interaction interface between the protein and G-quadruplex (D). Finally the structural parameters of the G-quadruplex are given by the angles between the guanines for each tetrad (E-G).

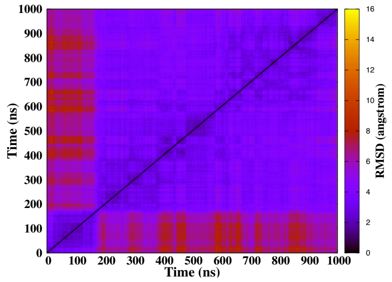


(A)

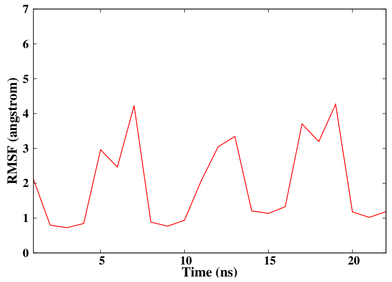


(B)

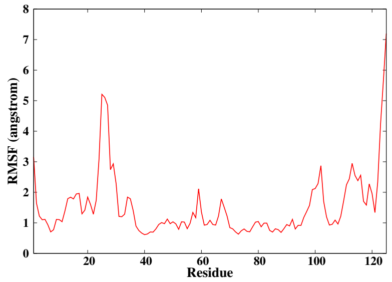


(C)

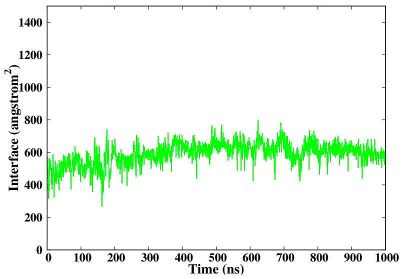


(D)

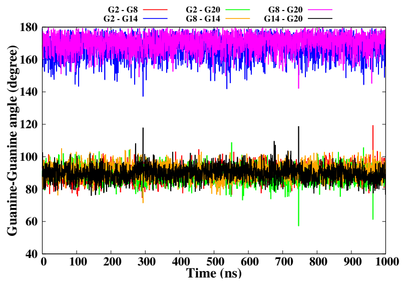


(E)

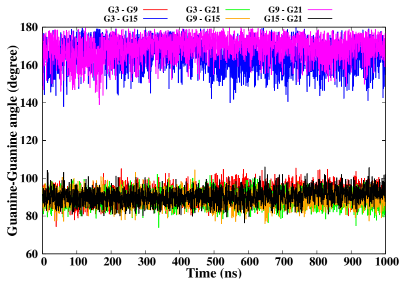


(F)

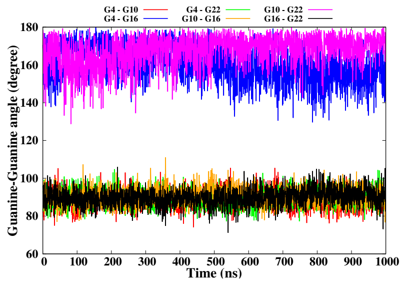


(G)

**Figure S5** – Simulation of the h-Telo G-quadruplex DNA in interaction with 2E4 according to the model 8-4, run 1. The convergence of the simulation is given by the RMSD-2D map of the DNA-Protein complex (A). The mobility of the DNA and protein residues is given by their root mean square fluctuation (B-C). Surface of the interaction interface between the protein and G-quadruplex (D). Finally the structural parameters of the G-quadruplex are given by the angles between the guanines for each tetrad (E-G).

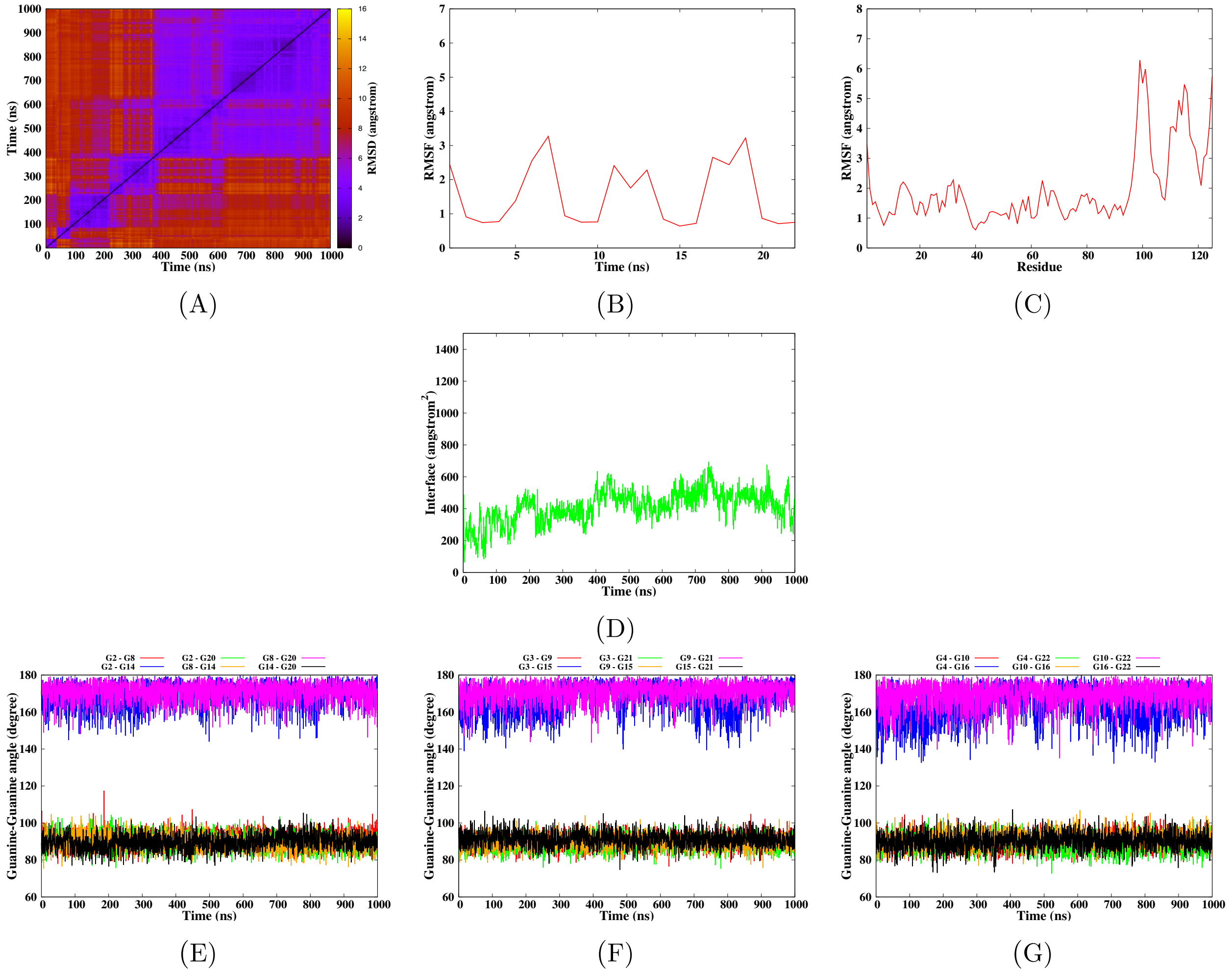


**Figure S6** – Simulation of the h-Telo G-quadruplex DNA in interaction with 2E4 according to the model 8-4, run 2. The convergence of the simulation is given by the RMSD-2D map of the DNA-Protein complex (A). The mobility of the DNA and protein residues is given by their root mean square fluctuation (B-C). Surface of the interaction interface between the protein and G-quadruplex (D). Finally the structural parameters of the G-quadruplex are given by the angles between the guanines for each tetrad (E-G).

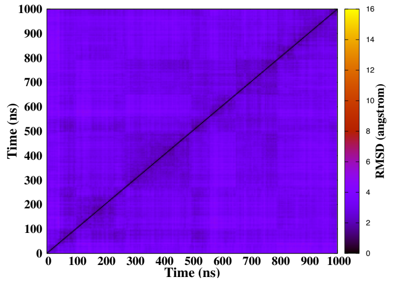


(A)

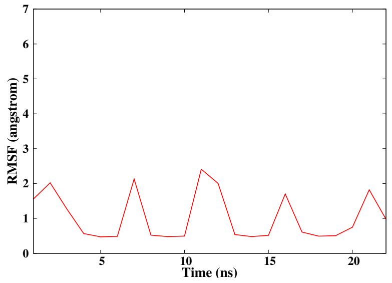


(B)

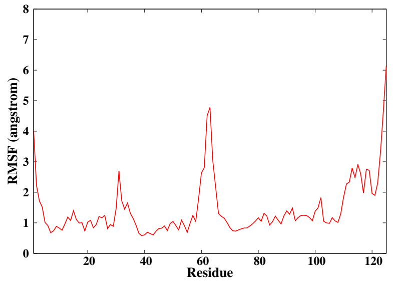


(C)

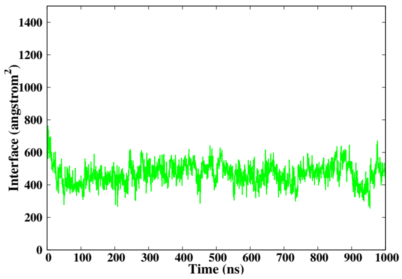


(D)

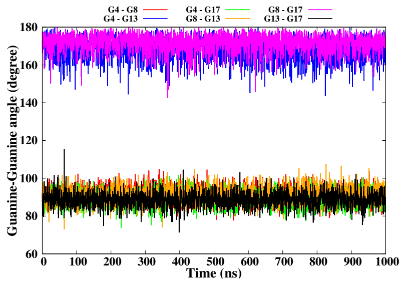


(E)

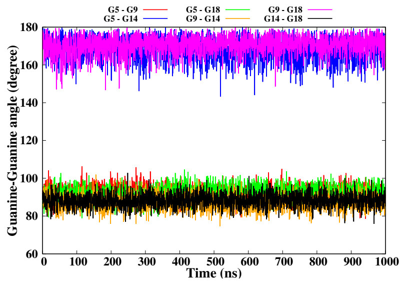


(F)

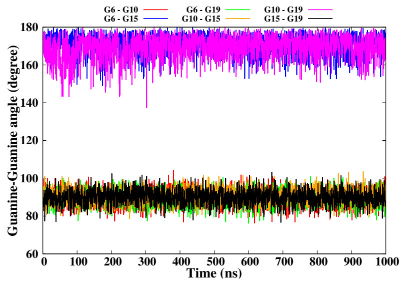


(G)

**Figure S7** – Simulation of the c-Myc G-quadruplex DNA in interaction with 2E4 according to the model 1-1, run 1. The convergence of the simulation is given by the RMSD-2D map of the DNA-Protein complex (A). The mobility of the DNA and protein residues is given by their root mean square fluctuation (B-C). Surface of the interaction interface between the protein and G-quadruplex (D). Finally the structural parameters of the G-quadruplex are given by the angles between the guanines for each tetrad (E-G).

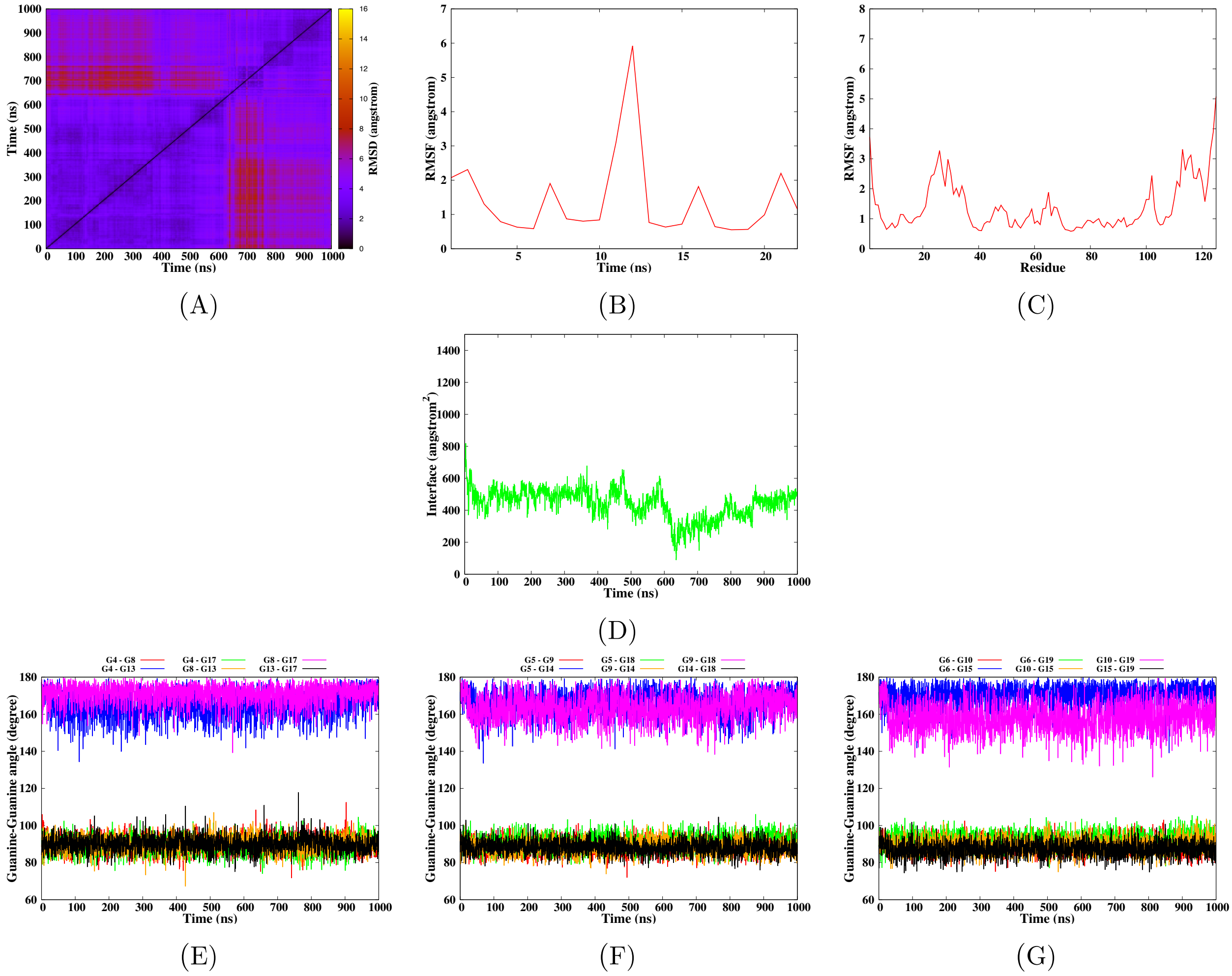


**Figure S8** – Simulation of the c-Myc G-quadruplex DNA in interaction with 2E4 according to the model 1-1, run 2. The convergence of the simulation is given by the RMSD-2D map of the DNA-Protein complex (A). The mobility of the DNA and protein residues is given by their root mean square fluctuation (B-C). Surface of the interaction interface between the protein and G-quadruplex (D). Finally the structural parameters of the G-quadruplex are given by the angles between the guanines for each tetrad (E-G).

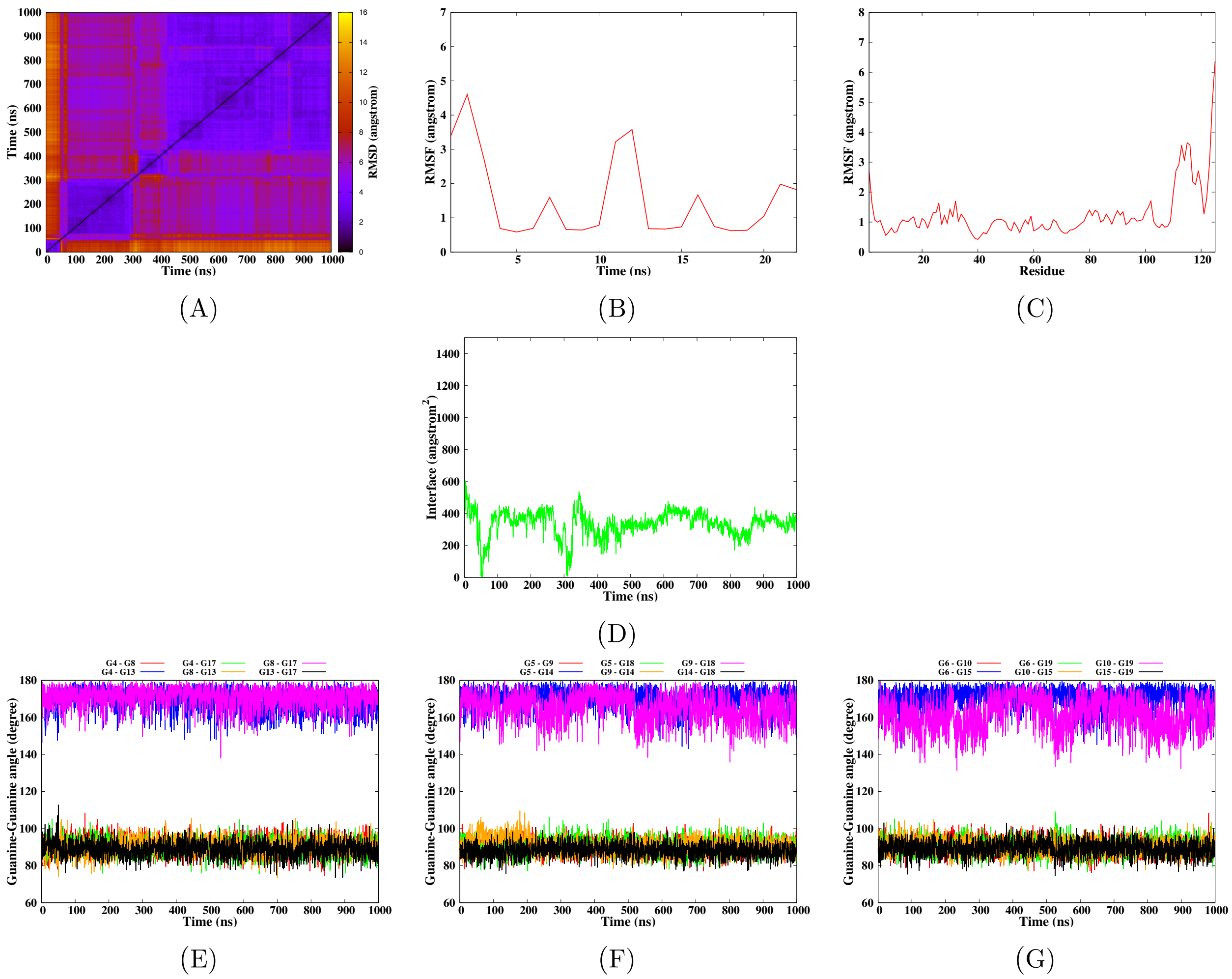


**Figure S9** – Simulation of the c-Myc G-quadruplex DNA in interaction with 2E4 according to the model 5-4, run 1. The convergence of the simulation is given by the RMSD-2D map of the DNA-Protein complex (A). The mobility of the DNA and protein residues is given by their root mean square fluctuation (B-C). Surface of the interaction interface between the protein and G-quadruplex (D). Finally the structural parameters of the G-quadruplex are given by the angles between the guanines for each tetrad (E-G).

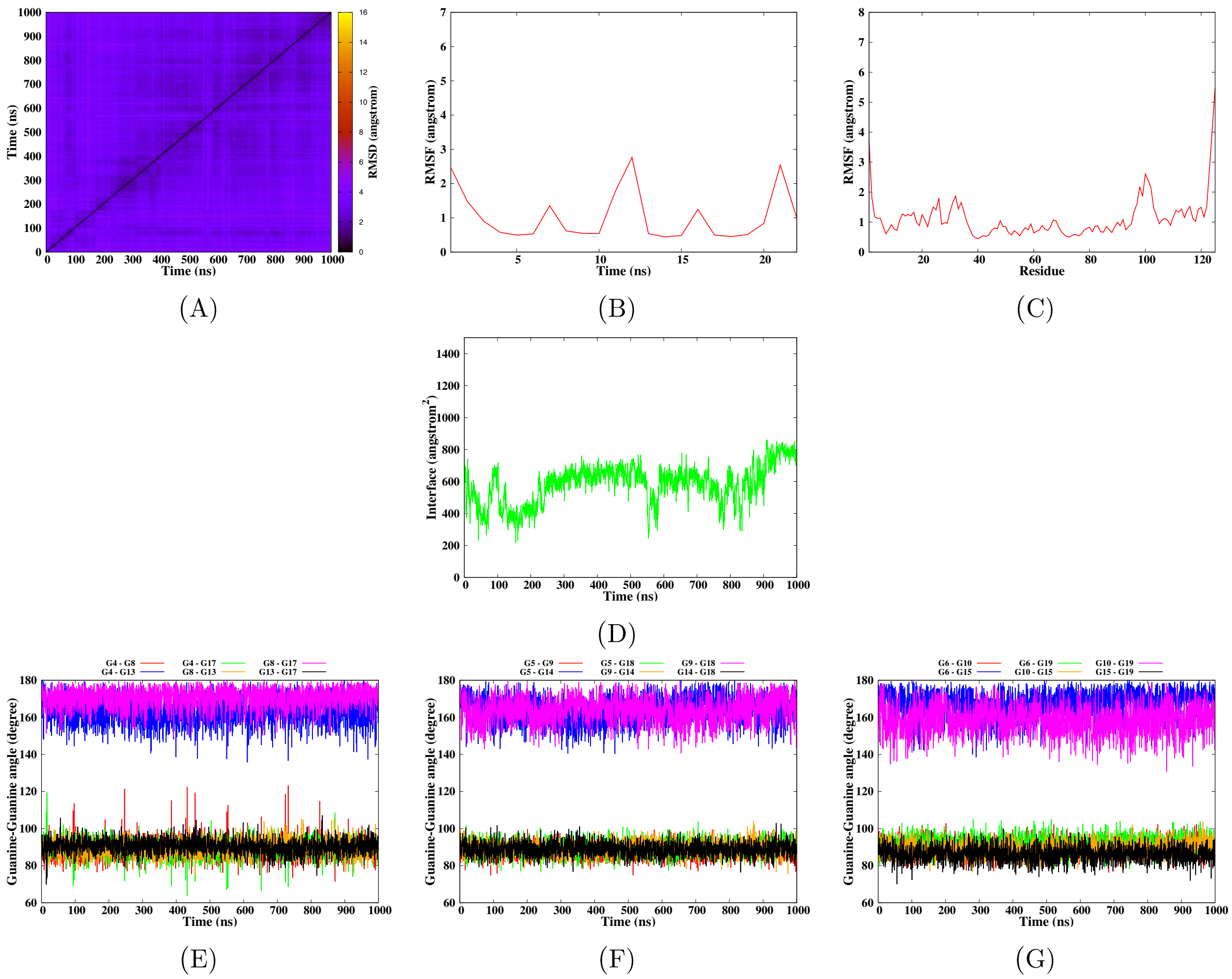


**Figure S10** – Simulation of the c-Myc G-quadruplex DNA in interaction with 2E4 according to the model 5-4, run 2. The convergence of the simulation is given by the RMSD-2D map of the DNA-Protein complex (A). The mobility of the DNA and protein residues is given by their root mean square fluctuation (B-C). Surface of the interaction interface between the protein and G-quadruplex (D). Finally the structural parameters of the G-quadruplex are given by the angles between the guanines for each tetrad (E-G).

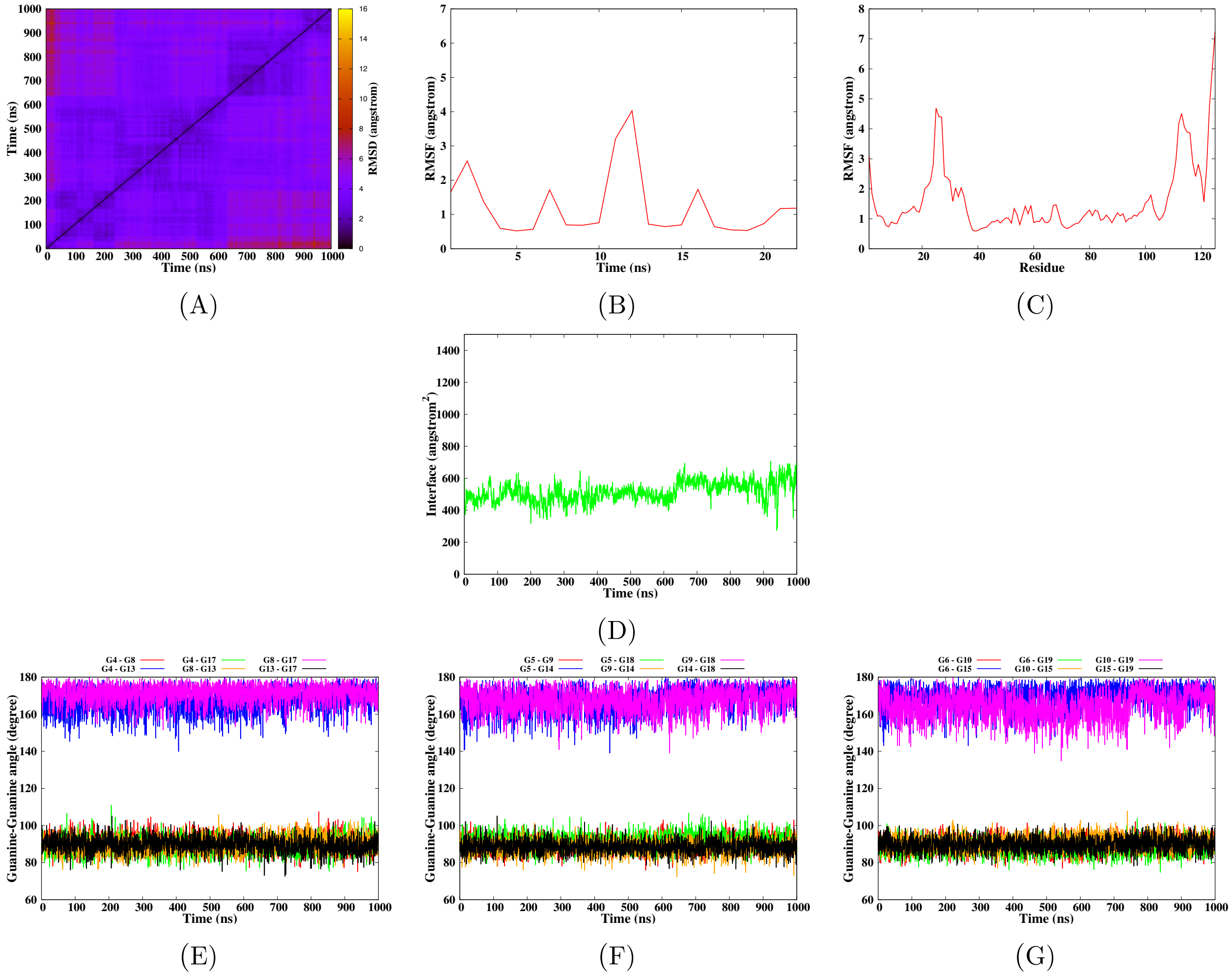


**Figure S11** – Simulation of the c-Myc G-quadruplex DNA in interaction with 2E4 according to the model 6-4, run 1. The convergence of the simulation is given by the RMSD-2D map of the DNA-Protein complex (A). The mobility of the DNA and protein residues is given by their root mean square fluctuation (B-C). Surface of the interaction interface between the protein and G-quadruplex (D). Finally the structural parameters of the G-quadruplex are given by the angles between the guanines for each tetrad (E-G).

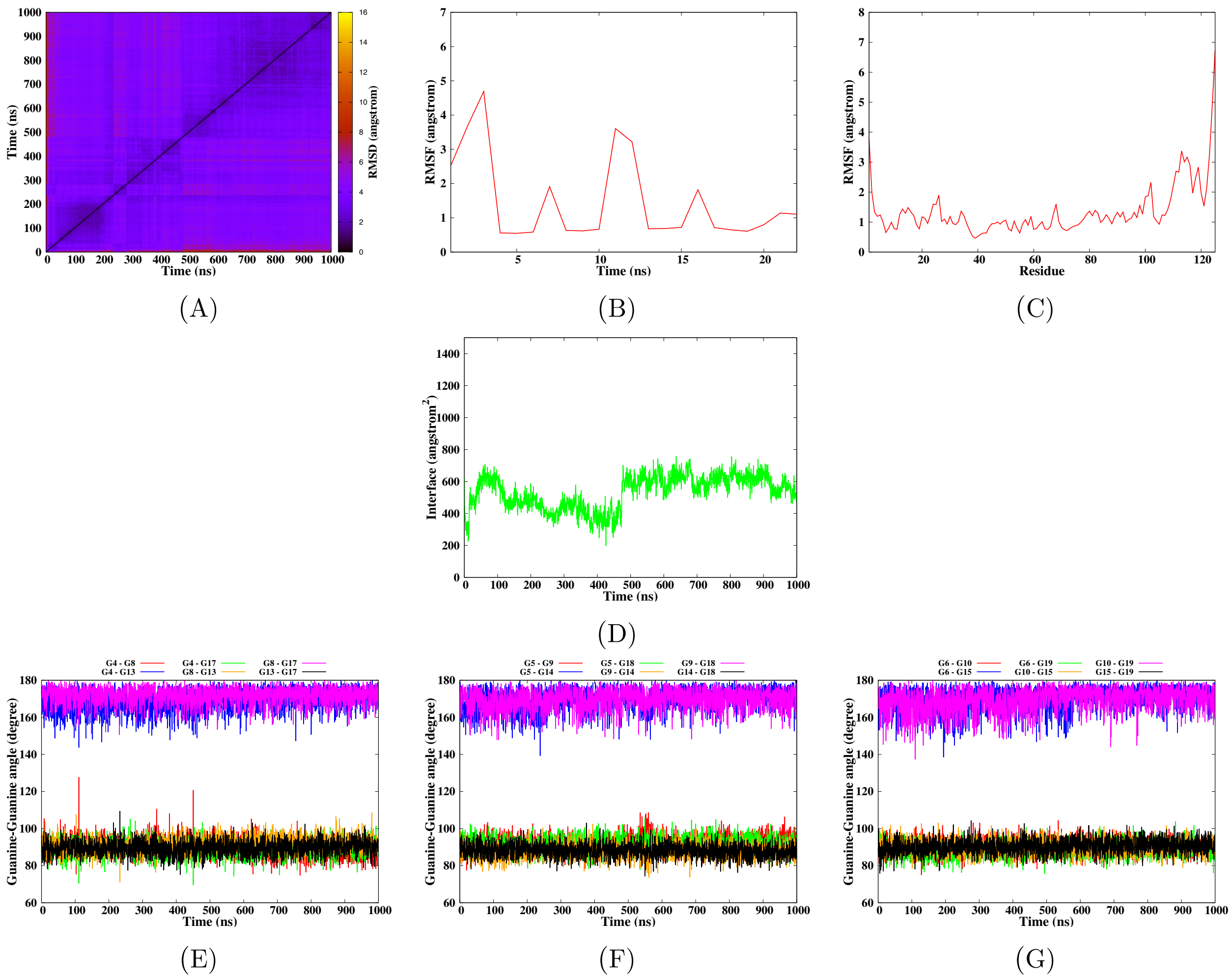


**Figure S12** – Simulation of the c-Myc G-quadruplex DNA in interaction with 2E4 according to the model 6-4, run 2. The convergence of the simulation is given by the RMSD-2D map of the DNA-Protein complex (A). The mobility of the DNA and protein residues is given by their root mean square fluctuation (B-C). Surface of the interaction interface between the protein and G-quadruplex (D). Finally the structural parameters of the G-quadruplex are given by the angles between the guanines for each tetrad (E-G).

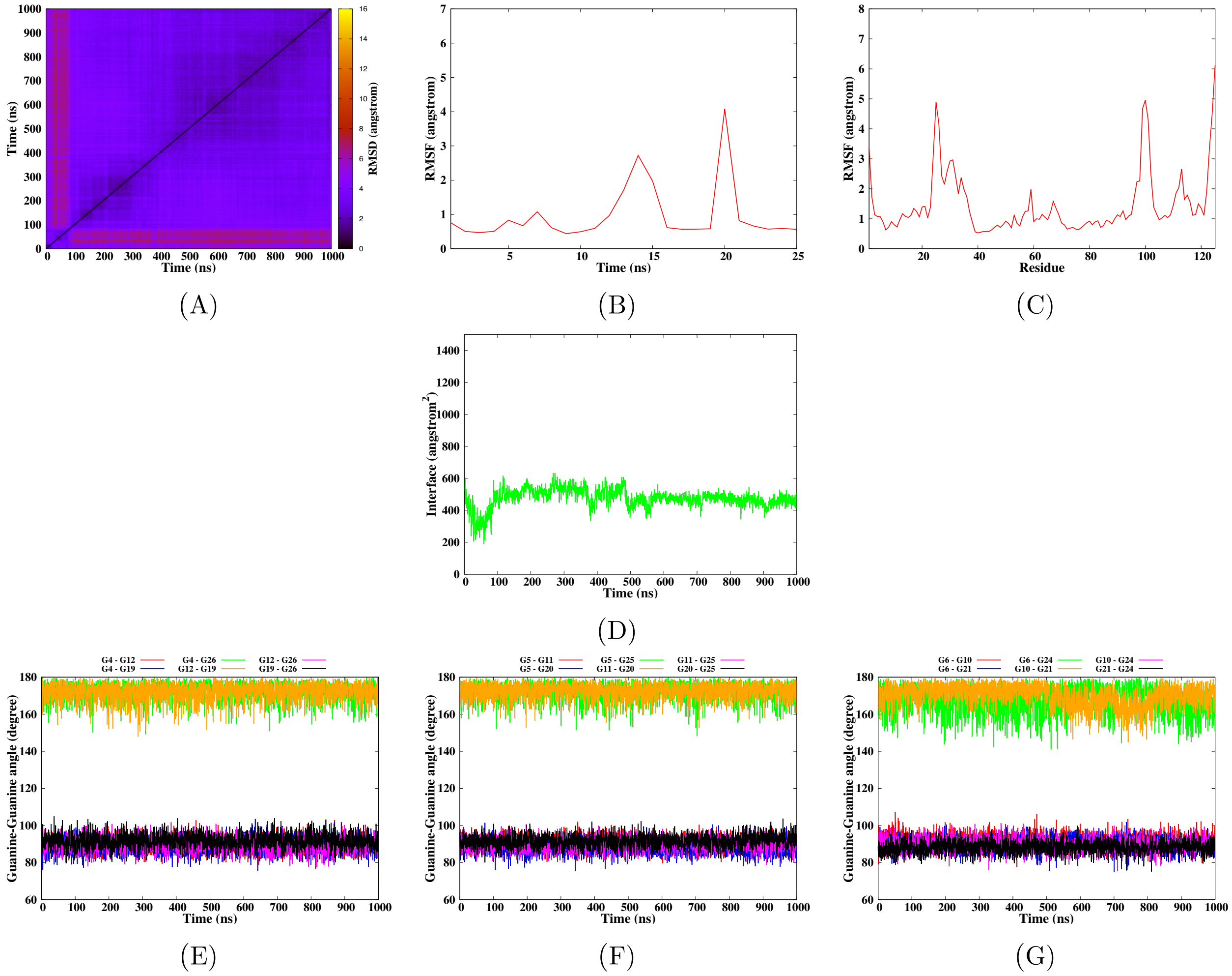


**Figure S13** – Simulation of the Bcl-2 G-quadruplex DNA in interaction with 2E4 according to the model 1-1, run 1. The convergence of the simulation is given by the RMSD-2D map of the DNA-Protein complex (A). The mobility of the DNA and protein residues is given by their root mean square fluctuation (B-C). Surface of the interaction interface between the protein and G-quadruplex (D). Finally the structural parameters of the G-quadruplex are given by the angles between the guanines for each tetrad (E-G).

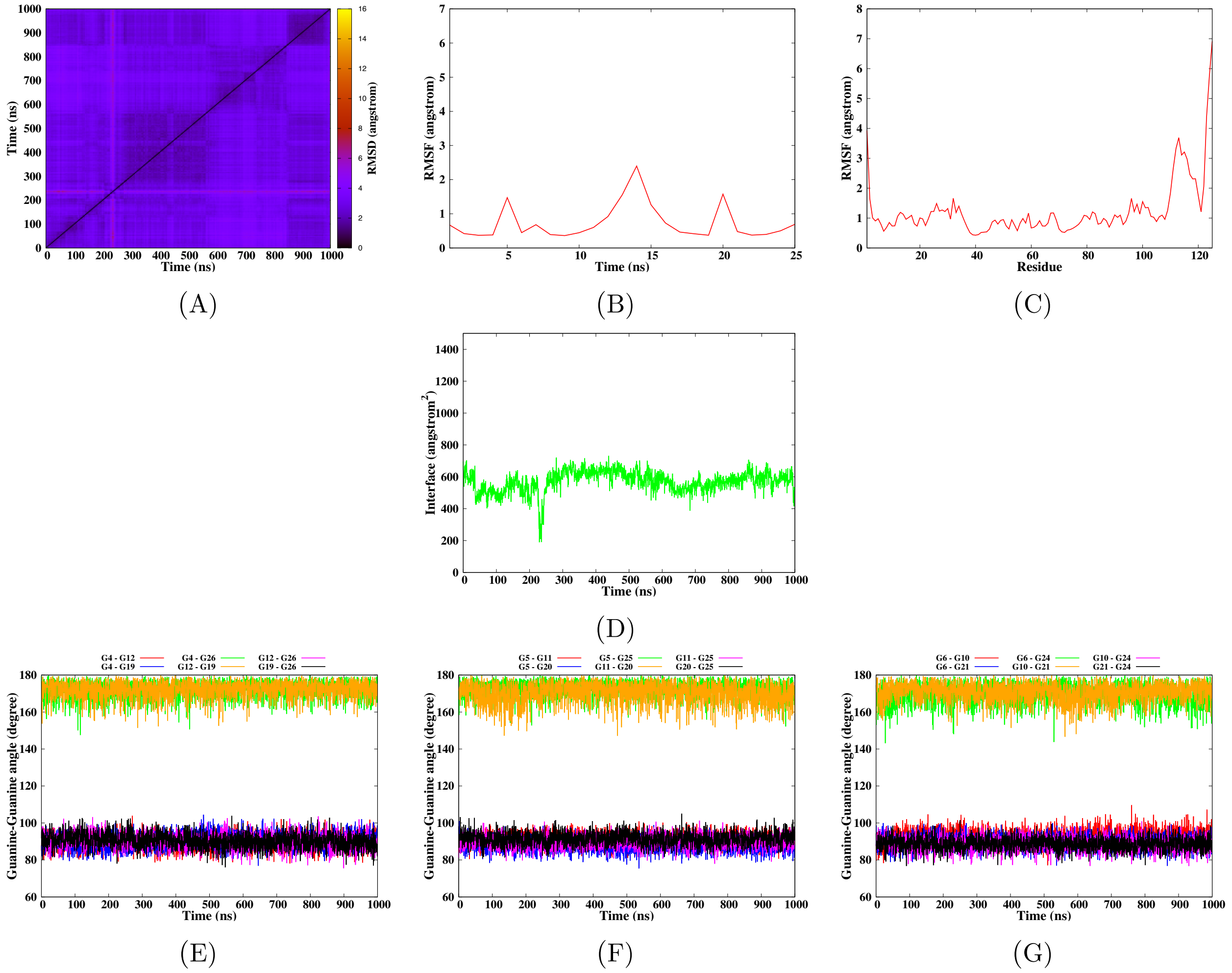


**Figure S14** – Simulation of the Bcl-2 G-quadruplex DNA in interaction with 2E4 according to the model 1-1, run 2. The convergence of the simulation is given by the RMSD-2D map of the DNA-Protein complex (A). The mobility of the DNA and protein residues is given by their root mean square fluctuation (B-C). Surface of the interaction interface between the protein and G-quadruplex (D). Finally the structural parameters of the G-quadruplex are given by the angles between the guanines for each tetrad (E-G).

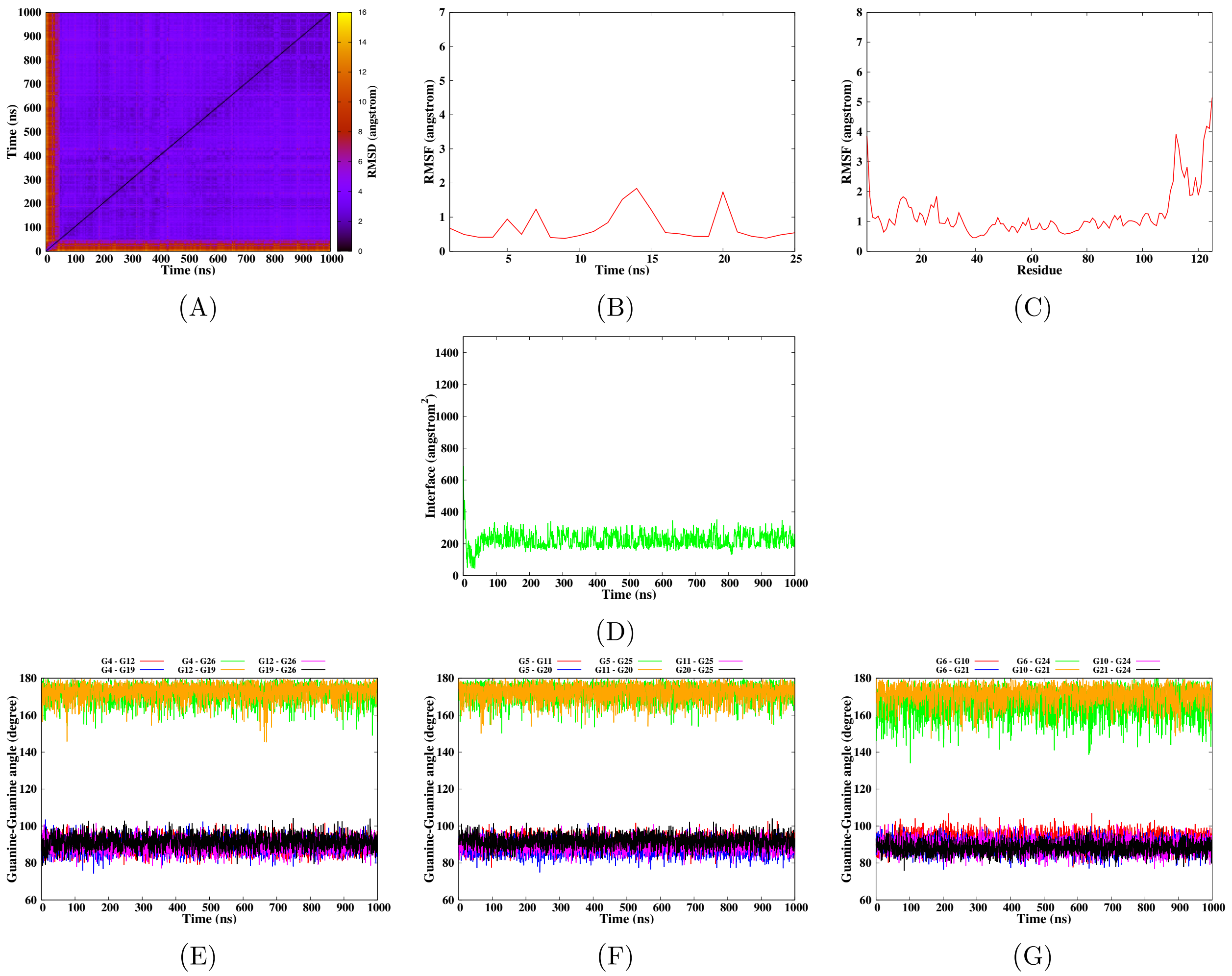


**Figure S15** – Simulation of the Bcl-2 G-quadruplex DNA in interaction with 2E4 according to the model 5-1, run 1. The convergence of the simulation is given by the RMSD-2D map of the DNA-Protein complex (A). The mobility of the DNA and protein residues is given by their root mean square fluctuation (B-C). Surface of the interaction interface between the protein and G-quadruplex (D). Finally the structural parameters of the G-quadruplex are given by the angles between the guanines for each tetrad (E-G).

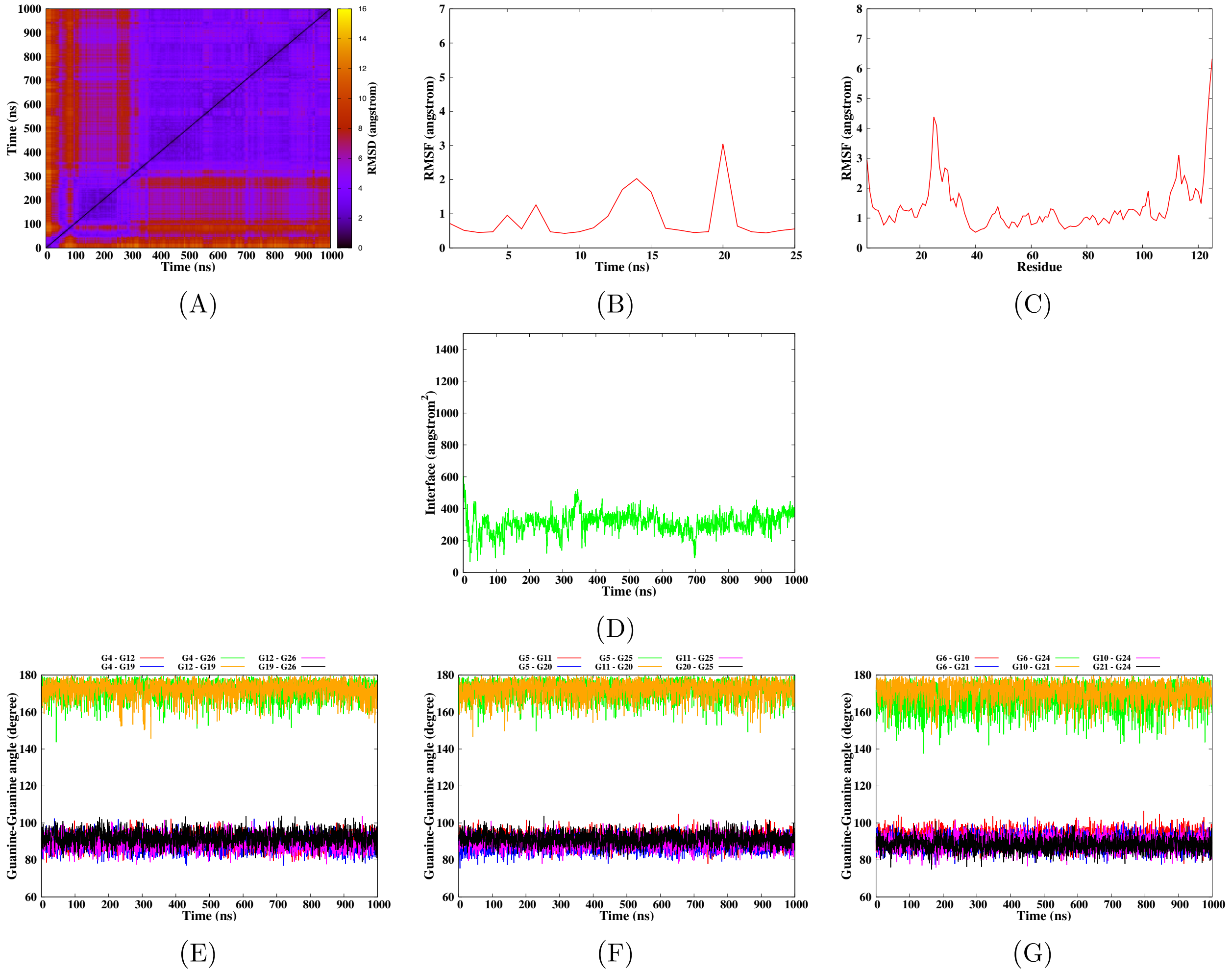


**Figure S16** – Simulation of the Bcl-2 G-quadruplex DNA in interaction with 2E4 according to the model 5-1, run 2. The convergence of the simulation is given by the RMSD-2D map of the DNA-Protein complex (A). The mobility of the DNA and protein residues is given by their root mean square fluctuation (B-C). Surface of the interaction interface between the protein and G-quadruplex (D). Finally the structural parameters of the G-quadruplex are given by the angles between the guanines for each tetrad (E-G).

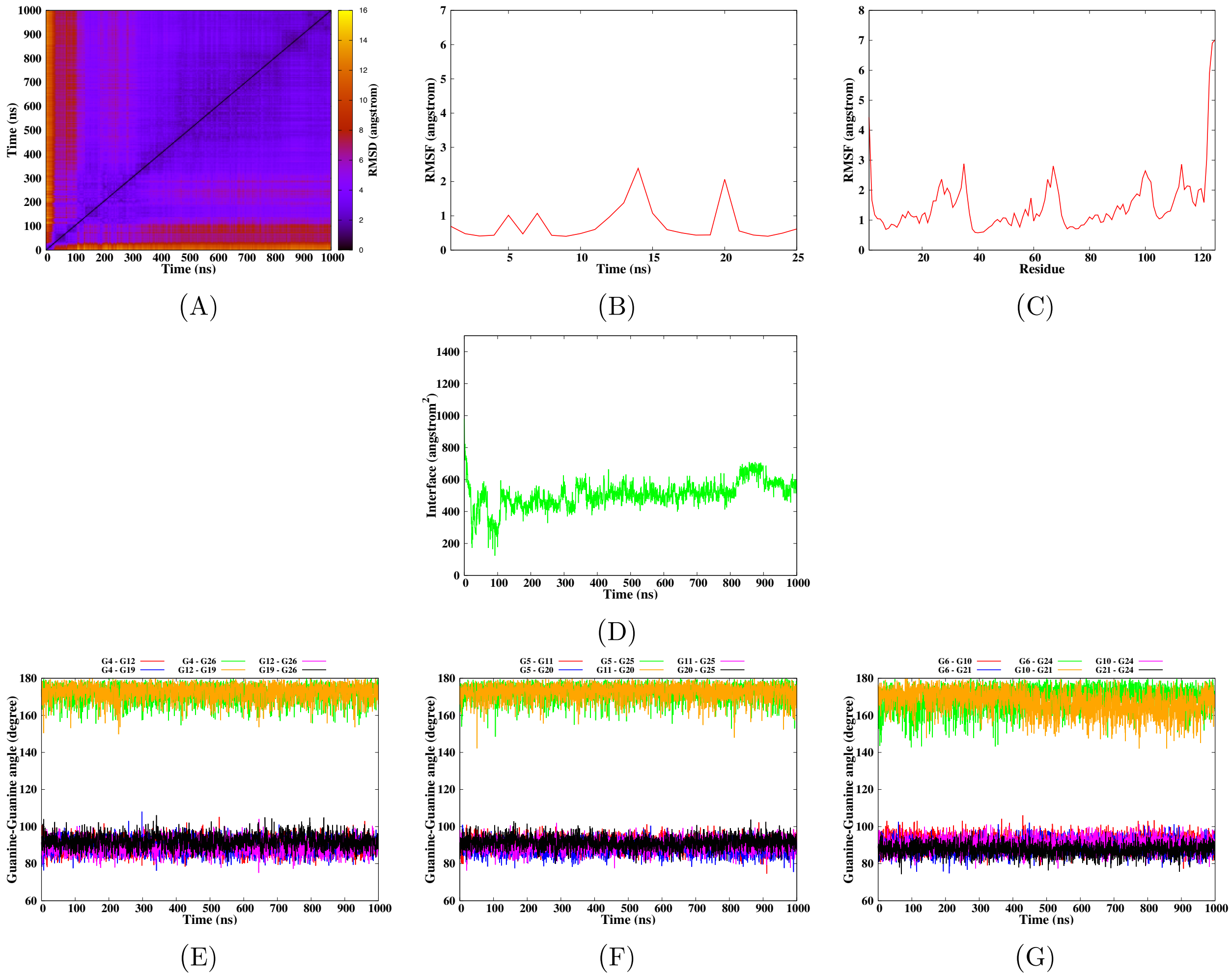


**Figure S17** – Simulation of the Bcl-2 G-quadruplex DNA in interaction with 2E4 according to the model 7-1, run 1. The convergence of the simulation is given by the RMSD-2D map of the DNA-Protein complex (A). The mobility of the DNA and protein residues is given by their root mean square fluctuation (B-C). Surface of the interaction interface between the protein and G-quadruplex (D). Finally the structural parameters of the G-quadruplex are given by the angles between the guanines for each tetrad (E-G).

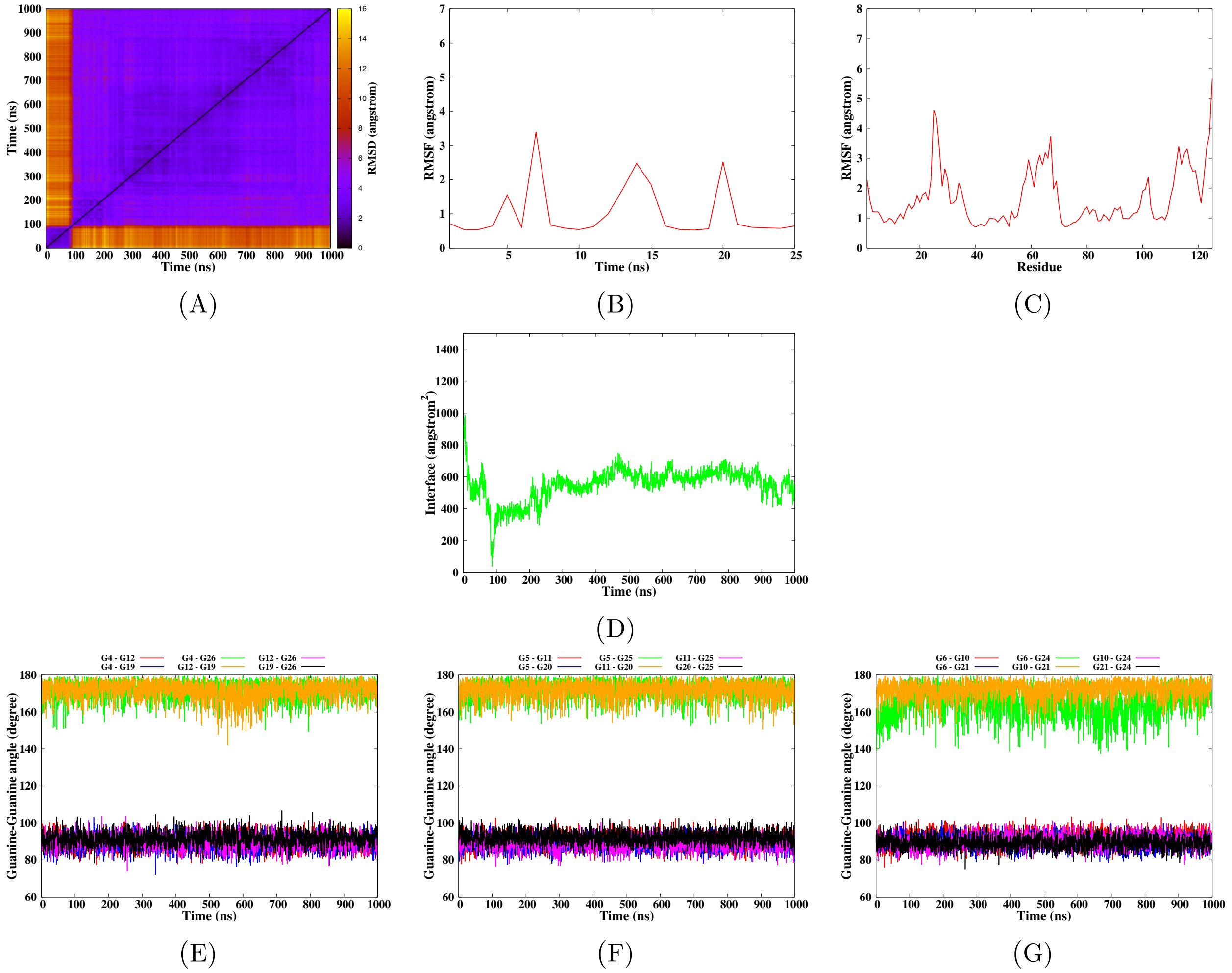


**Figure S18** – Simulation of the Bcl-2 G-quadruplex DNA in interaction with 2E4 according to the model 7-1, run 2. The convergence of the simulation is given by the RMSD-2D map of the DNA-Protein complex (A). The mobility of the DNA and protein residues is given by their root mean square fluctuation (B-C). Surface of the interaction interface between the protein and G-quadruplex (D). Finally the structural parameters of the G-quadruplex are given by the angles between the guanines for each tetrad (E-G).

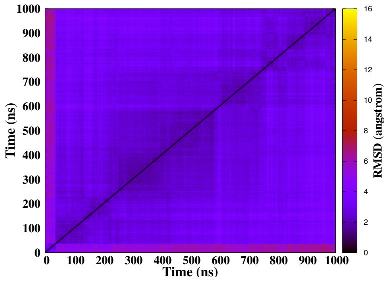


(A)

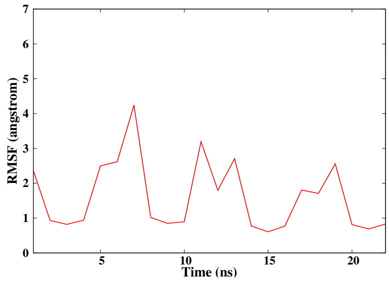


(B)

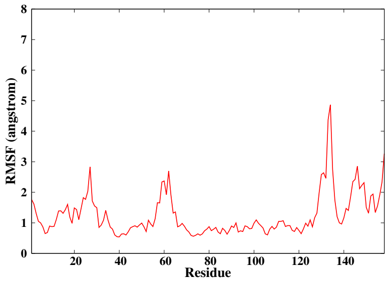


(C)

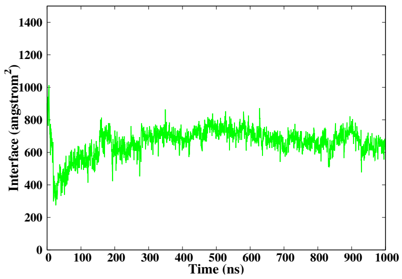


(D)

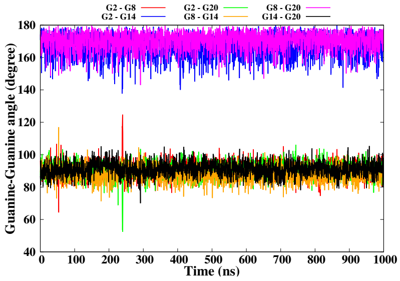


(E)

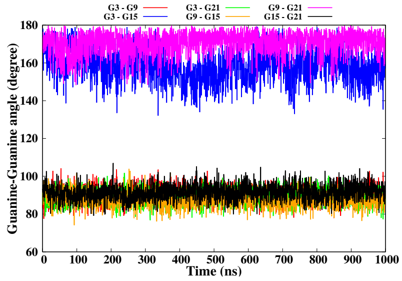


(F)

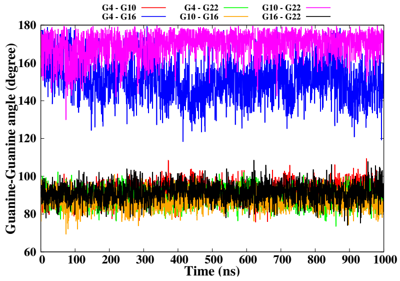


(G)

**Figure S19** – Simulation of the h-Telo G-quadruplex DNA in interaction with 2G10 according to the model 13-1, run 1. The convergence of the simulation is given by the RMSD-2D map of the DNA-Protein complex (A). The mobility of the DNA and protein residues is given by their root mean square fluctuation (B-C). Surface of the interaction interface between the protein and G-quadruplex (D). Finally the structural parameters of the G-quadruplex are given by the angles between the guanines for each tetrad (E-G).

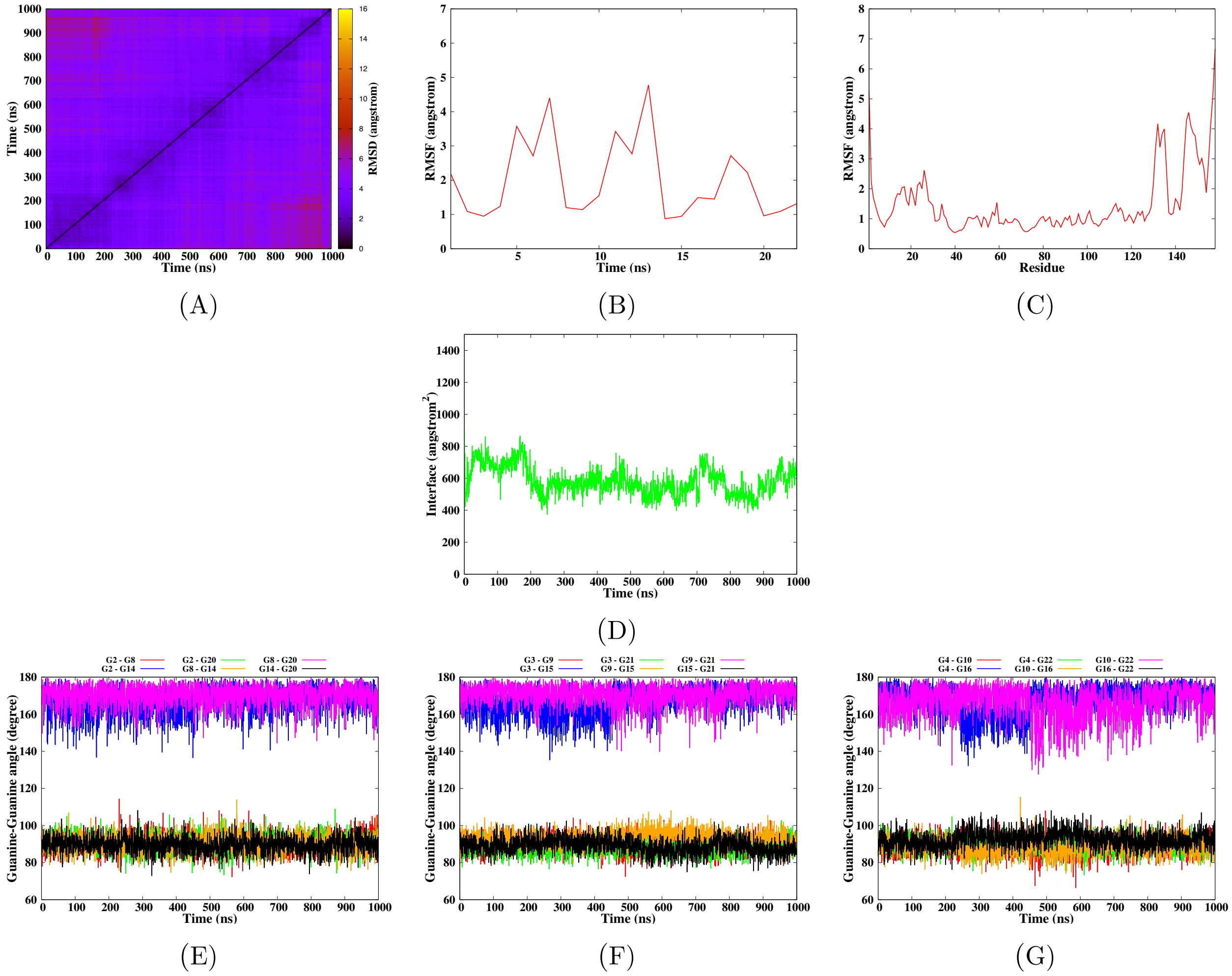


**Figure S20** – Simulation of the h-Telo G-quadruplex DNA in interaction with 2G10 according to the model 13-1, run 2. The convergence of the simulation is given by the RMSD-2D map of the DNA-Protein complex (A). The mobility of the DNA and protein residues is given by their root mean square fluctuation (B-C). Surface of the interaction interface between the protein and G-quadruplex (D). Finally the structural parameters of the G-quadruplex are given by the angles between the guanines for each tetrad (E-G).

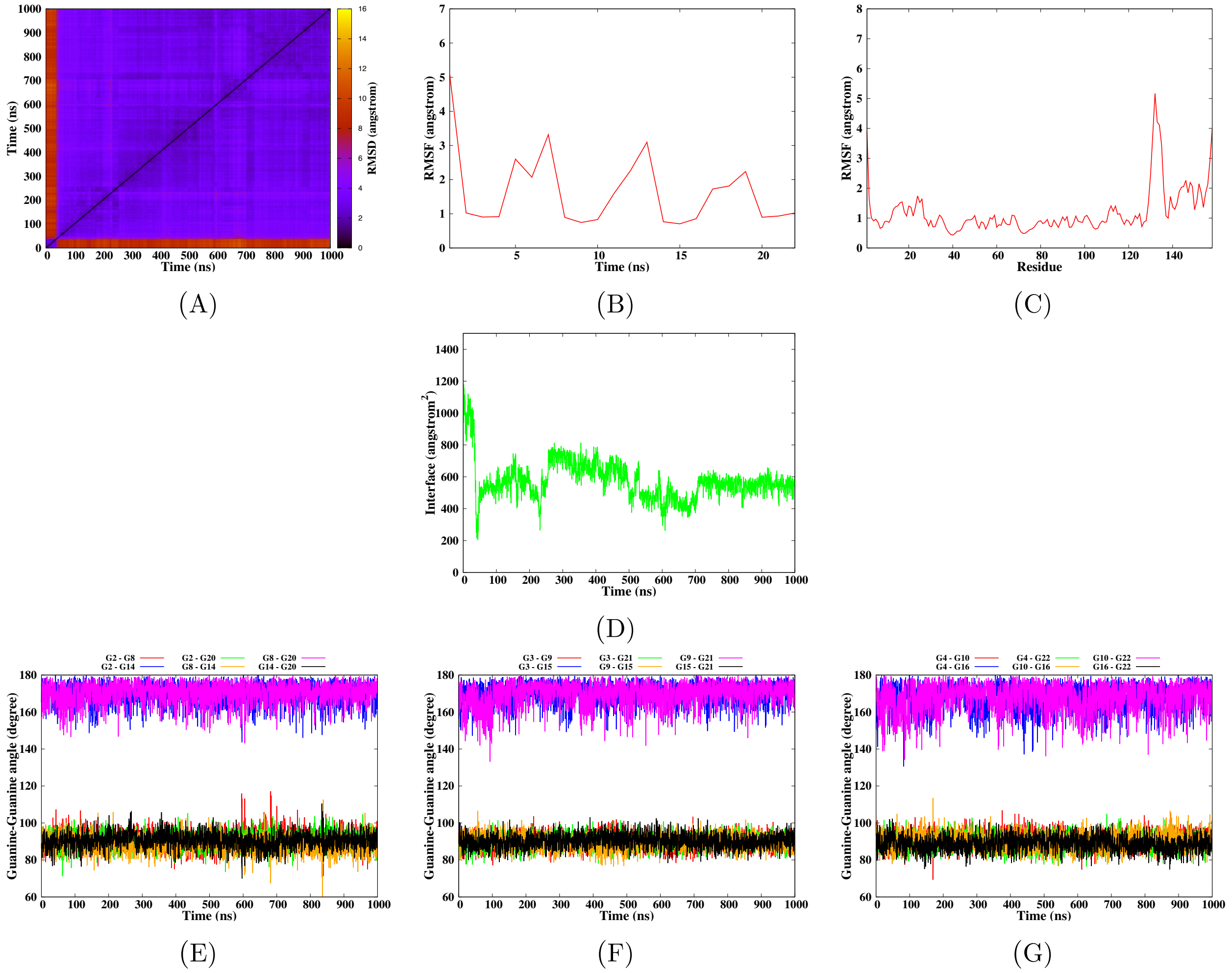


**Figure S21** – Simulation of the h-Telo G-quadruplex DNA in interaction with 2G10 according to the model 5-1, run 1. The convergence of the simulation is given by the RMSD-2D map of the DNA-Protein complex (A). The mobility of the DNA and protein residues is given by their root mean square fluctuation (B-C). Surface of the interaction interface between the protein and G-quadruplex (D). Finally the structural parameters of the G-quadruplex are given by the angles between the guanines for each tetrad (E-G).

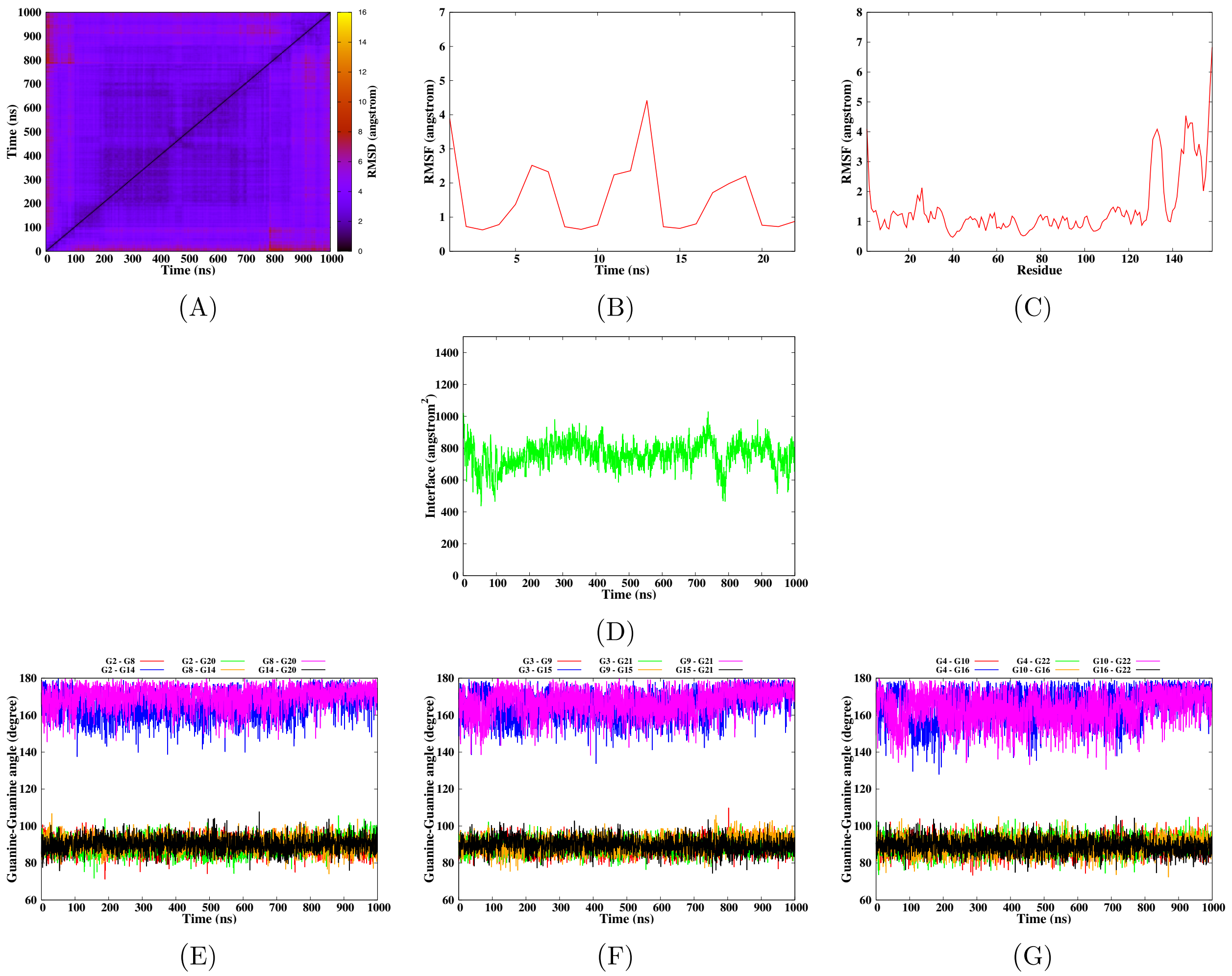

**Figure S22** – Simulation of the h-Telo G-quadruplex DNA in interaction with 2G10 according to the model 5-1, run 2. The convergence of the simulation is given by the RMSD-2D map of the DNA-Protein complex (A). The mobility of the DNA and protein residues is given by their root mean square fluctuation (B-C). Surface of the interaction interface between the protein and G-quadruplex (D). Finally the structural parameters of the G-quadruplex are given by the angles between the guanines for each tetrad (E-G).

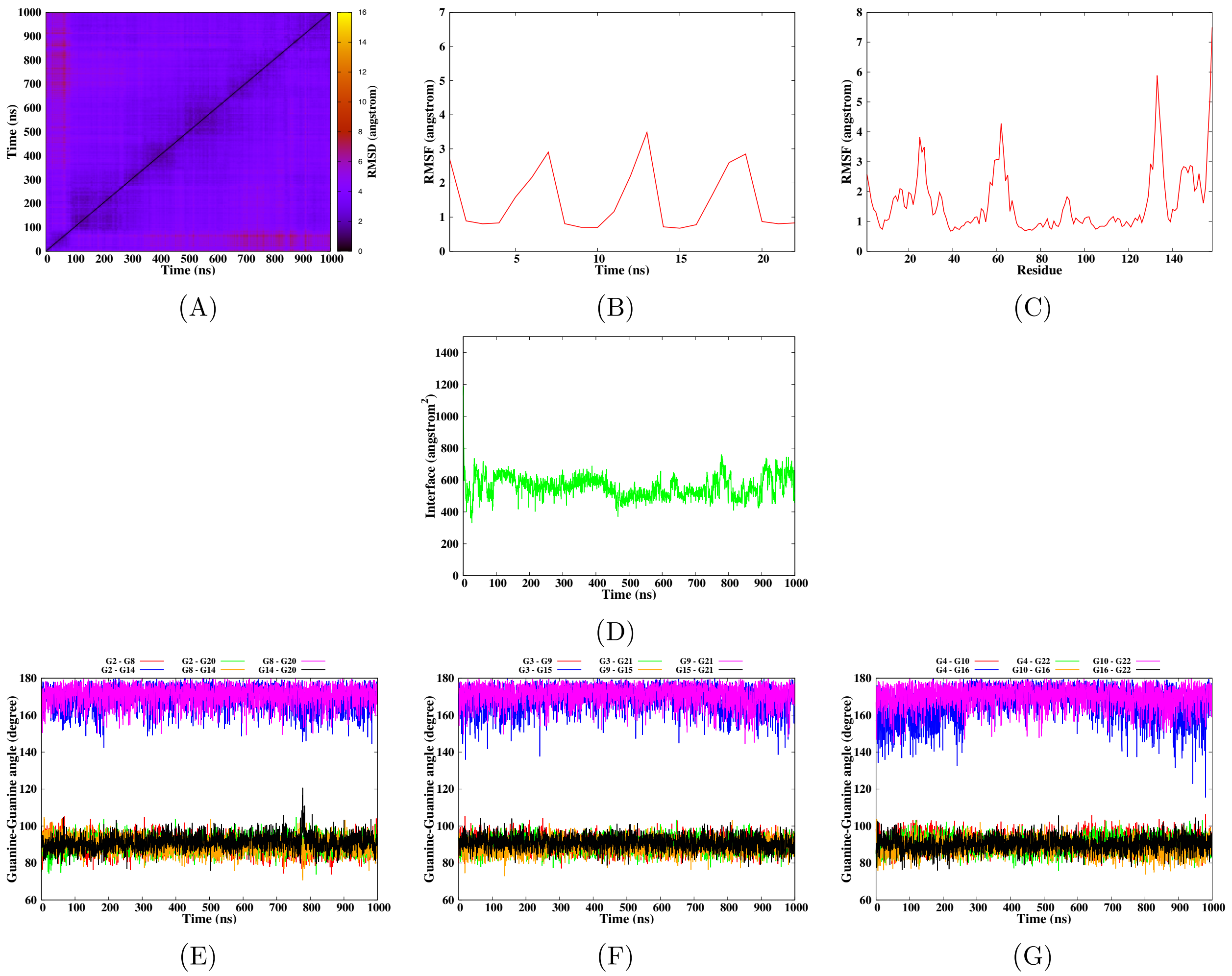


**Figure S23** – Simulation of the h-Telo G-quadruplex DNA in interaction with 2G10 according to the model 6-1, run 1. The convergence of the simulation is given by the RMSD-2D map of the DNA-Protein complex (A). The mobility of the DNA and protein residues is given by their root mean square fluctuation (B-C). Surface of the interaction interface between the protein and G-quadruplex (D). Finally the structural parameters of the G-quadruplex are given by the angles between the guanines for each tetrad (E-G).

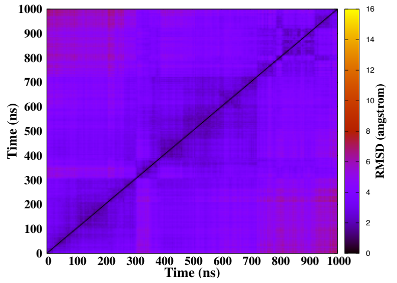


(A)

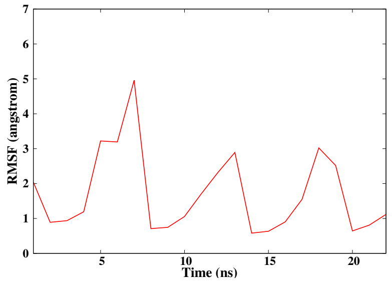


(B)

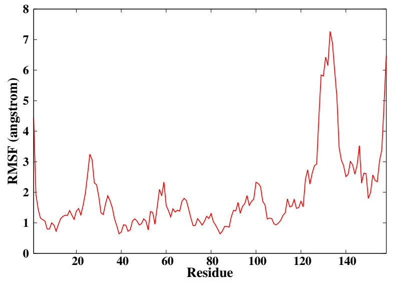


(C)

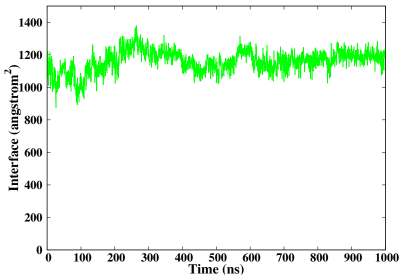


(D)

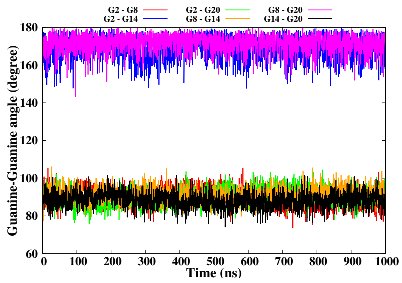


(E)

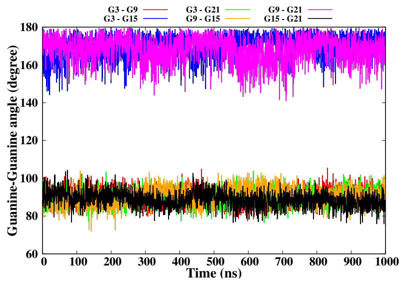


(F)

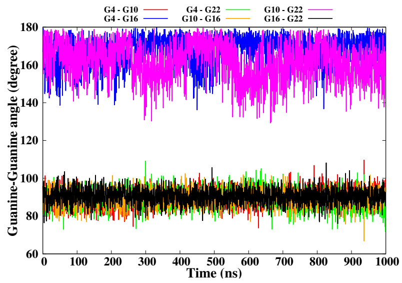


(G)

**Figure S24** – Simulation of the h-Telo G-quadruplex DNA in interaction with 2G10 according to the model 6-1, run 2. The convergence of the simulation is given by the RMSD-2D map of the DNA-Protein complex (A). The mobility of the DNA and protein residues is given by their root mean square fluctuation (B-C). Surface of the interaction interface between the protein and G-quadruplex (D). Finally the structural parameters of the G-quadruplex are given by the angles between the guanines for each tetrad (E-G).

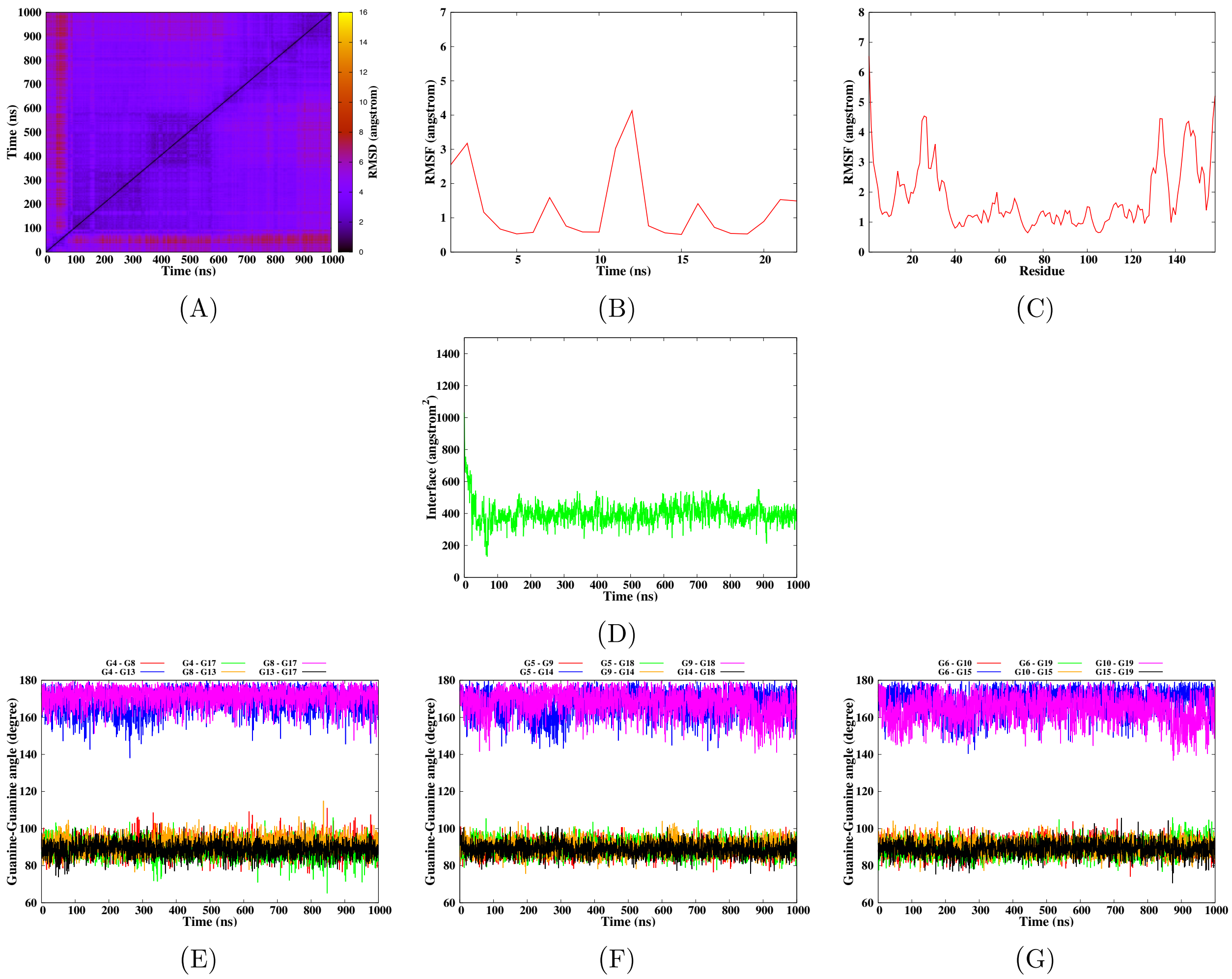


**Figure S25** – Simulation of the c-Myc G-quadruplex DNA in interaction with 2G10 according to the model 1-1, run 1. The convergence of the simulation is given by the RMSD-2D map of the DNA-Protein complex (A). The mobility of the DNA and protein residues is given by their root mean square fluctuation (B-C). Surface of the interaction interface between the protein and G-quadruplex (D). Finally the structural parameters of the G-quadruplex are given by the angles between the guanines for each tetrad (E-G).

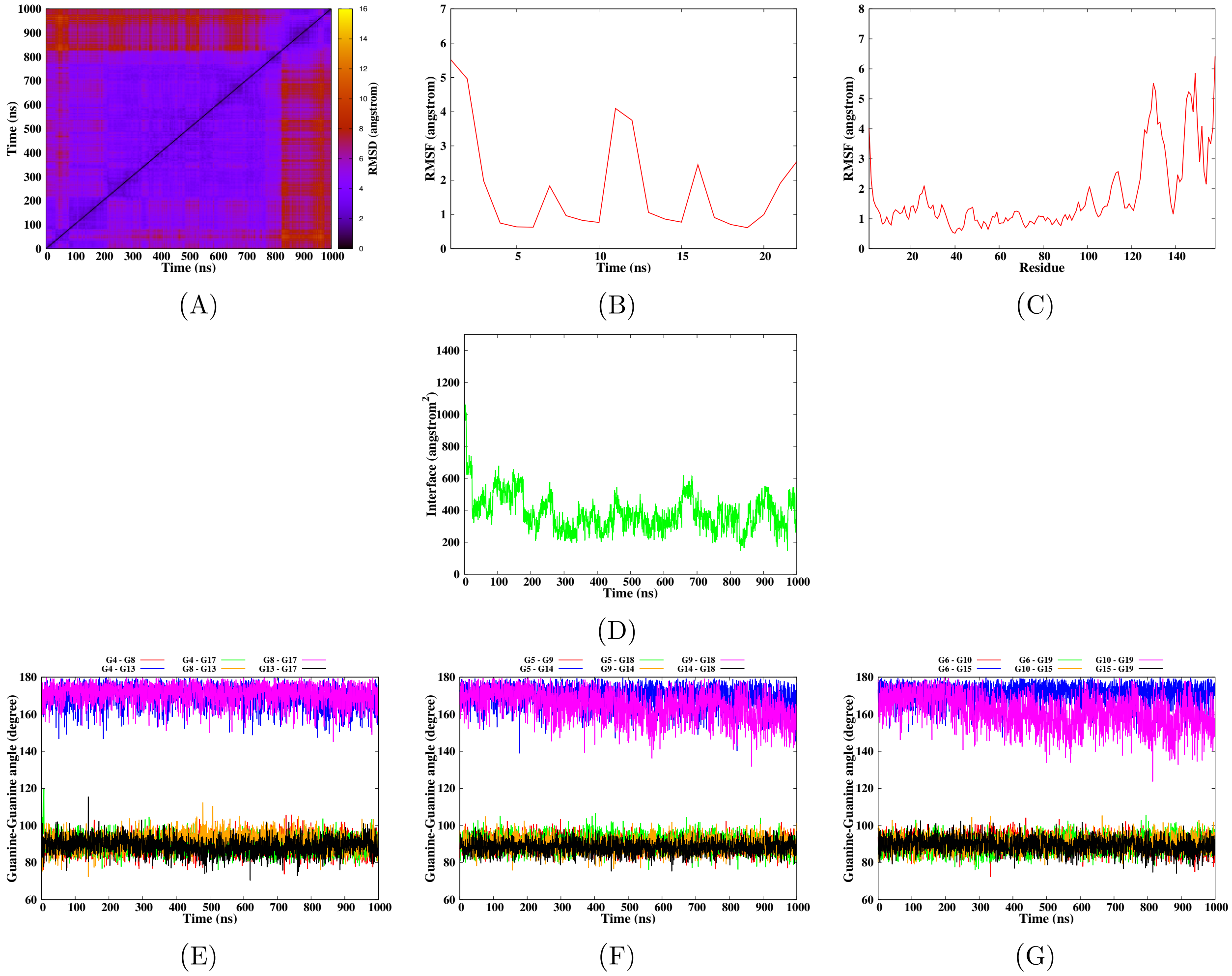


**Figure S26** – Simulation of the c-Myc G-quadruplex DNA in interaction with 2G10 according to the model 1-1, run 2. The convergence of the simulation is given by the RMSD-2D map of the DNA-Protein complex (A). The mobility of the DNA and protein residues is given by their root mean square fluctuation (B-C). Surface of the interaction interface between the protein and G-quadruplex (D). Finally the structural parameters of the G-quadruplex are given by the angles between the guanines for each tetrad (E-G).

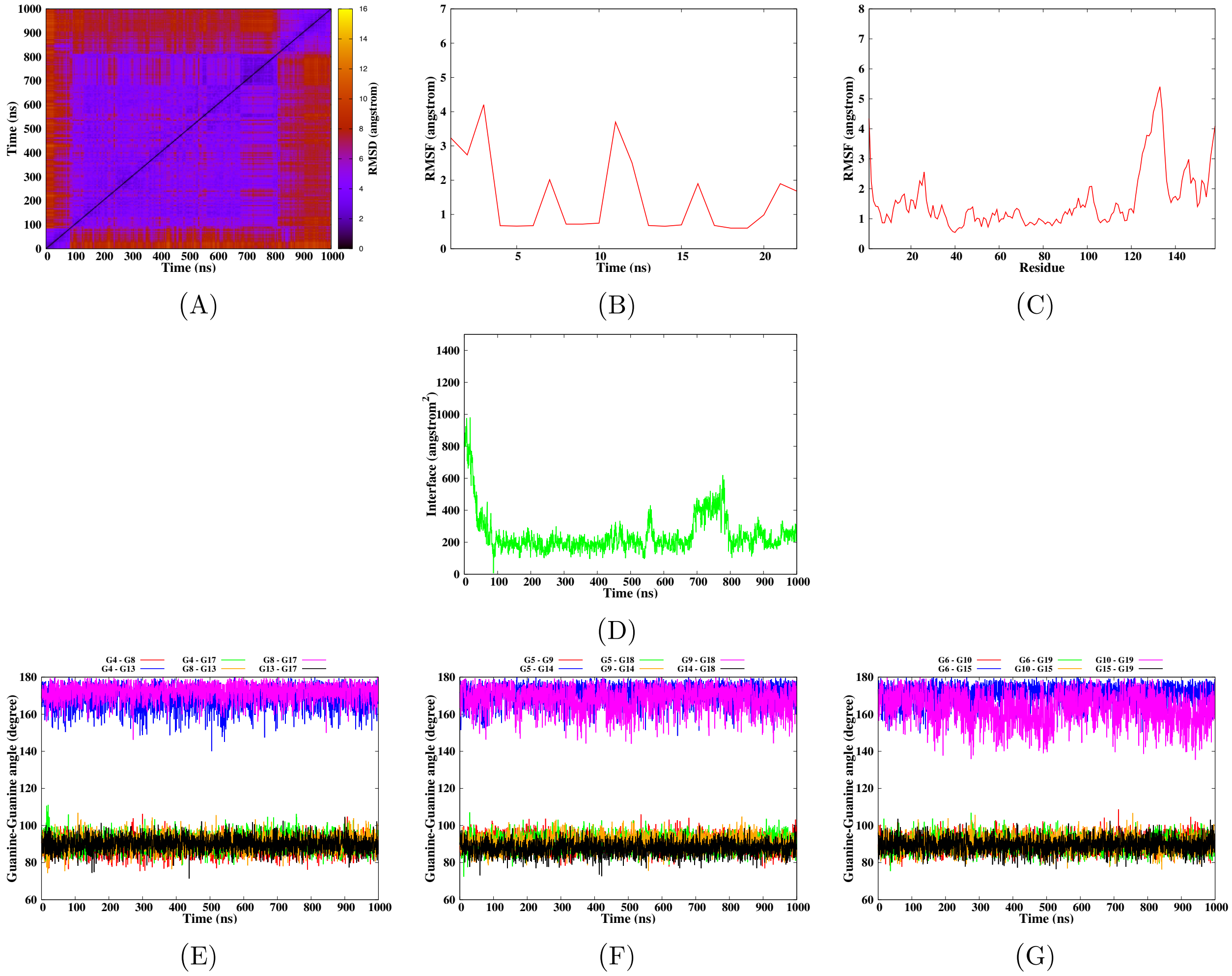


**Figure S27** – Simulation of the c-Myc G-quadruplex DNA in interaction with 2G10 according to the model 2-1, run 1. The convergence of the simulation is given by the RMSD-2D map of the DNA-Protein complex (A). The mobility of the DNA and protein residues is given by their root mean square fluctuation (B-C). Surface of the interaction interface between the protein and G-quadruplex (D). Finally the structural parameters of the G-quadruplex are given by the angles between the guanines for each tetrad (E-G).

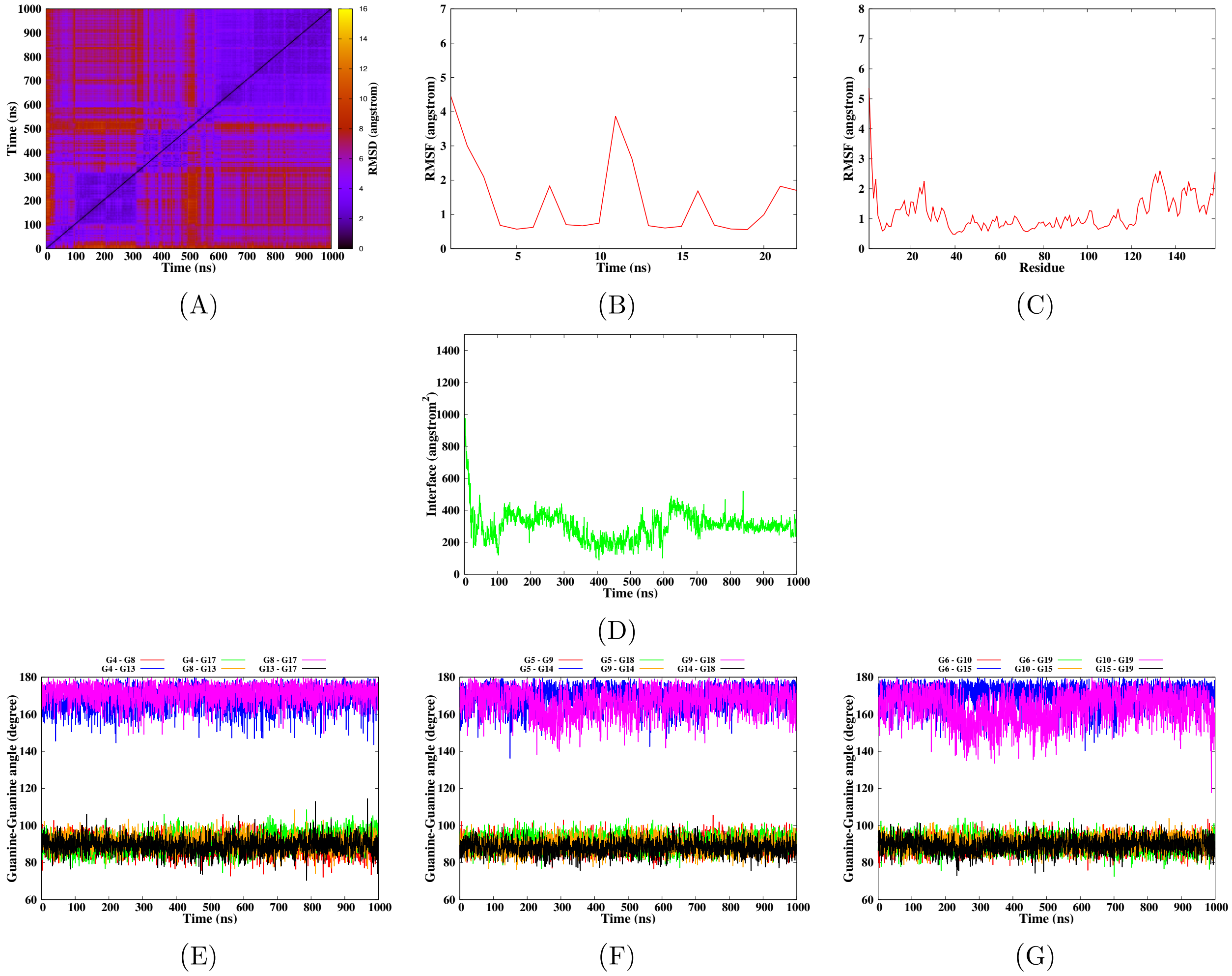


**Figure S28** – Simulation of the c-Myc G-quadruplex DNA in interaction with 2G10 according to the model 2-1, run 2. The convergence of the simulation is given by the RMSD-2D map of the DNA-Protein complex (A). The mobility of the DNA and protein residues is given by their root mean square fluctuation (B-C). Surface of the interaction interface between the protein and G-quadruplex (D). Finally the structural parameters of the G-quadruplex are given by the angles between the guanines for each tetrad (E-G).

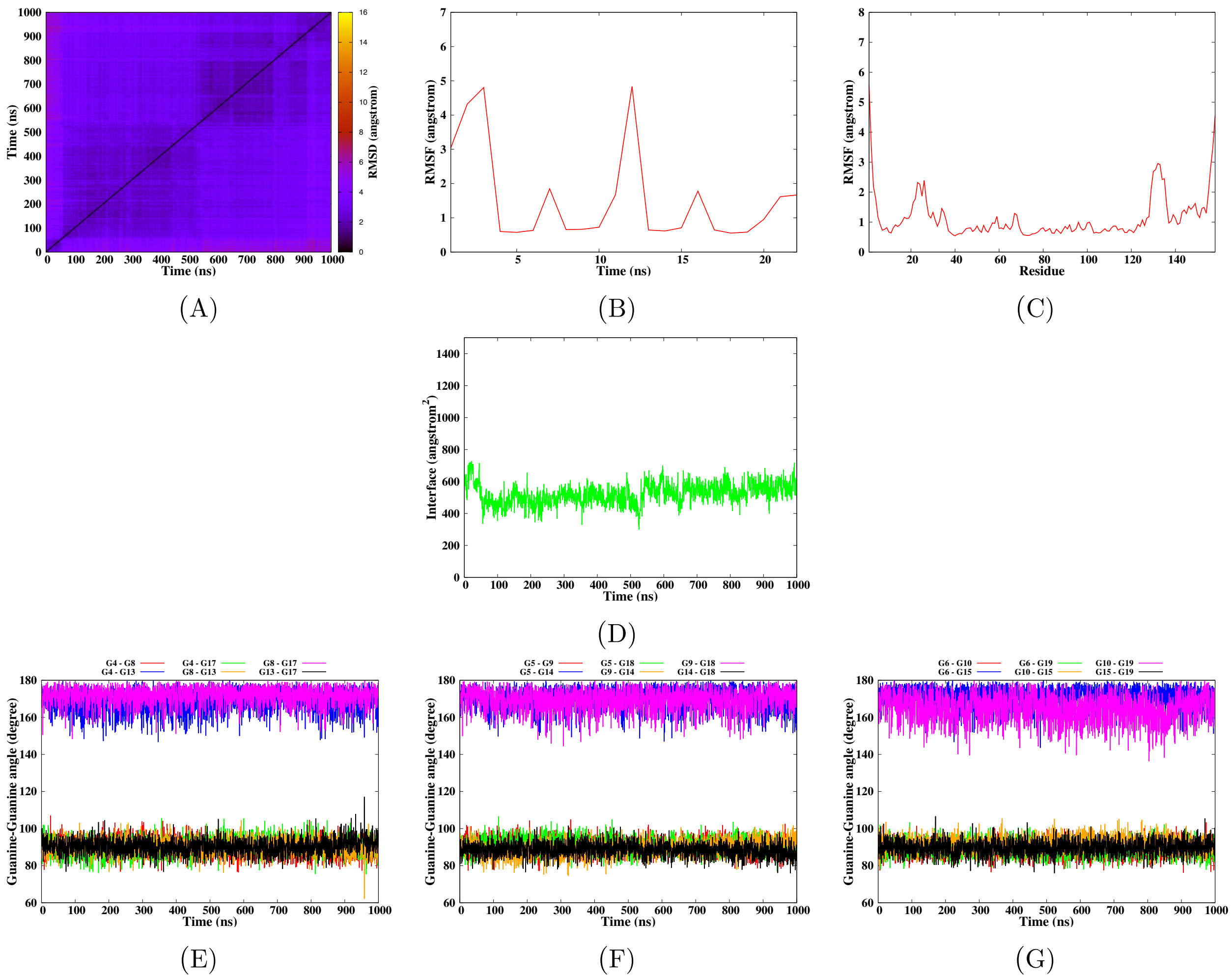


**Figure S29** – Simulation of the c-Myc G-quadruplex DNA in interaction with 2G10 according to the model 3-1, run 1. The convergence of the simulation is given by the RMSD-2D map of the DNA-Protein complex (A). The mobility of the DNA and protein residues is given by their root mean square fluctuation (B-C). Surface of the interaction interface between the protein and G-quadruplex (D). Finally the structural parameters of the G-quadruplex are given by the angles between the guanines for each tetrad (E-G).

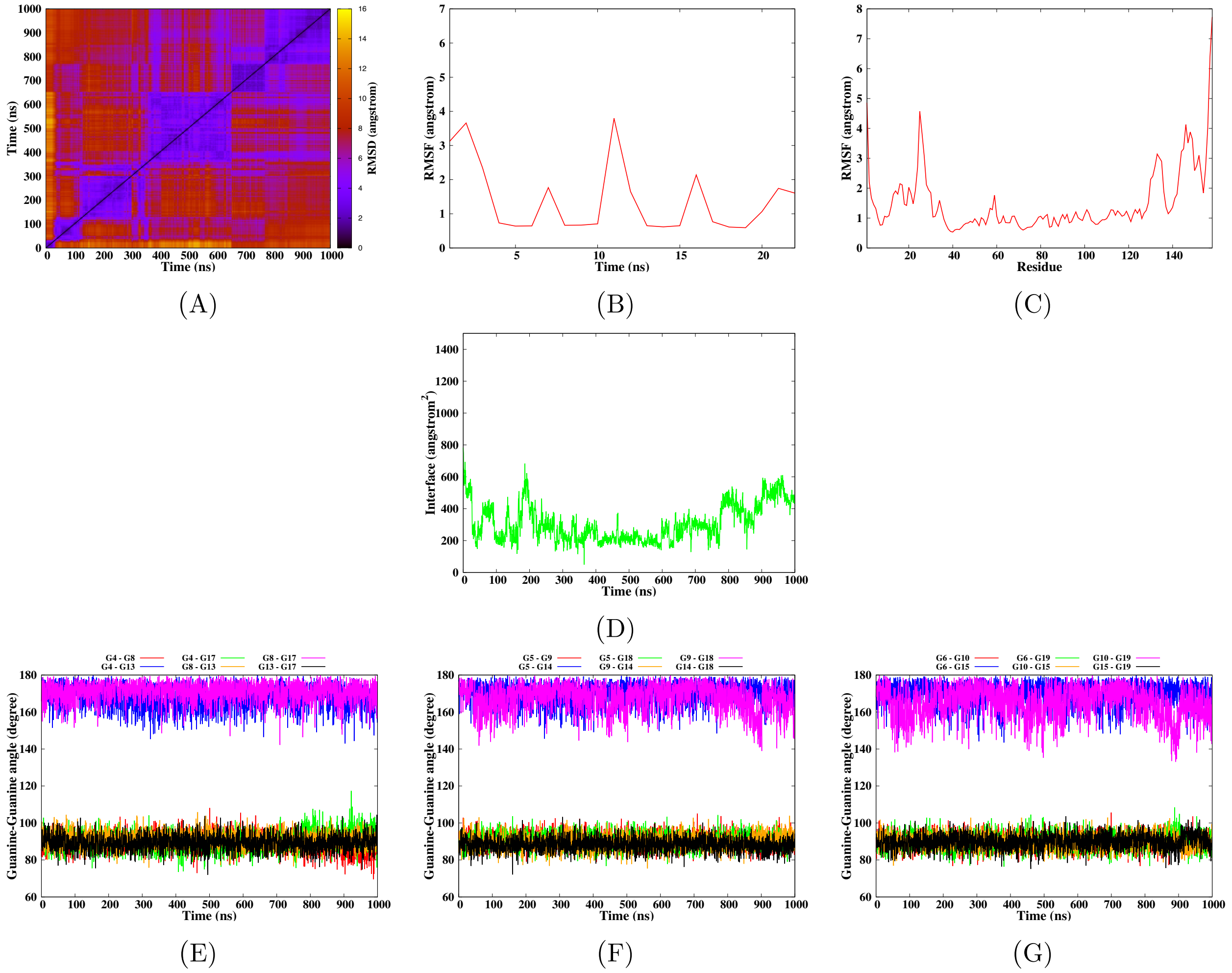


**Figure S30** – Simulation of the c-Myc G-quadruplex DNA in interaction with 2G10 according to the model 3-1, run 2. The convergence of the simulation is given by the RMSD-2D map of the DNA-Protein complex (A). The mobility of the DNA and protein residues is given by their root mean square fluctuation (B-C). Surface of the interaction interface between the protein and G-quadruplex (D). Finally the structural parameters of the G-quadruplex are given by the angles between the guanines for each tetrad (E-G).

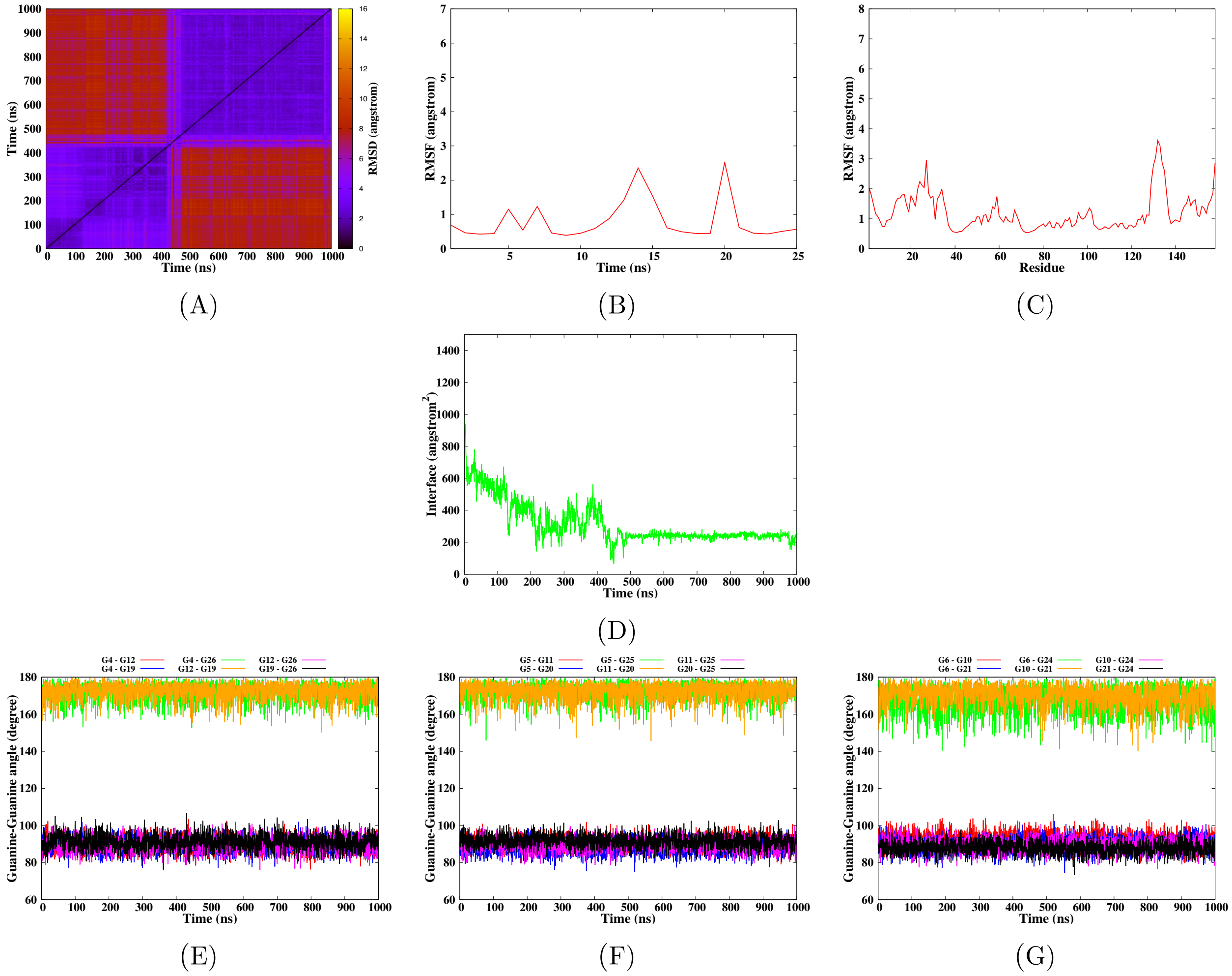


**Figure S31** – Simulation of the Bcl-2 G-quadruplex DNA in interaction with 2G10 according to the model 1-1, run 1. The convergence of the simulation is given by the RMSD-2D map of the DNA-Protein complex (A). The mobility of the DNA and protein residues is given by their root mean square fluctuation (B-C). Surface of the interaction interface between the protein and G-quadruplex (D). Finally the structural parameters of the G-quadruplex are given by the angles between the guanines for each tetrad (E-G).

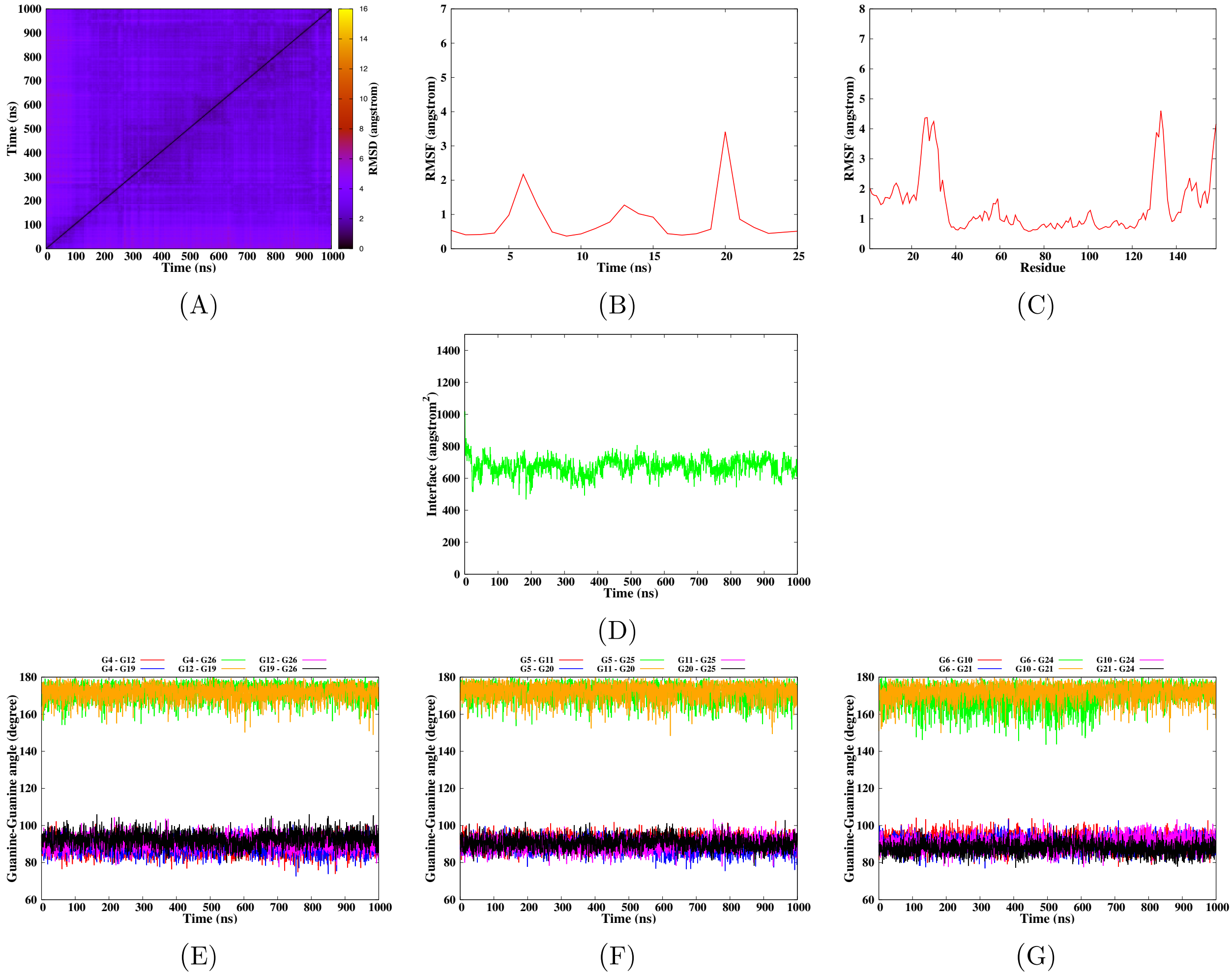


**Figure S32** – Simulation of the Bcl-2 G-quadruplex DNA in interaction with 2G10 according to the model 1-1, run 2. The convergence of the simulation is given by the RMSD-2D map of the DNA-Protein complex (A). The mobility of the DNA and protein residues is given by their root mean square fluctuation (B-C). Surface of the interaction interface between the protein and G-quadruplex (D). Finally the structural parameters of the G-quadruplex are given by the angles between the guanines for each tetrad (E-G).

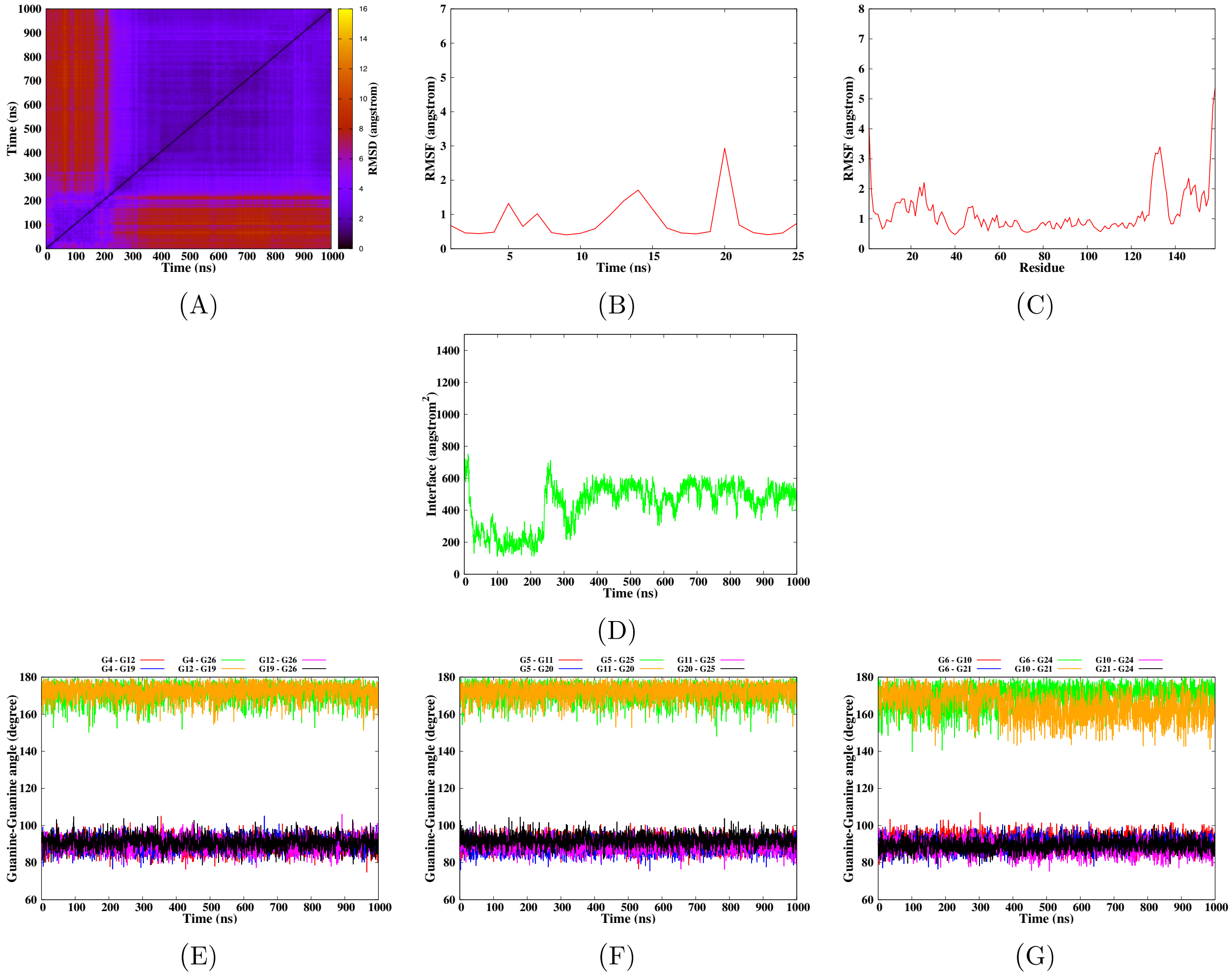


**Figure S33** – Simulation of the Bcl-2 G-quadruplex DNA in interaction with 2G10 according to the model 13-1, run 1. The convergence of the simulation is given by the RMSD-2D map of the DNA-Protein complex (A). The mobility of the DNA and protein residues is given by their root mean square fluctuation (B-C). Surface of the interaction interface between the protein and G-quadruplex (D). Finally the structural parameters of the G-quadruplex are given by the angles between the guanines for each tetrad (E-G).

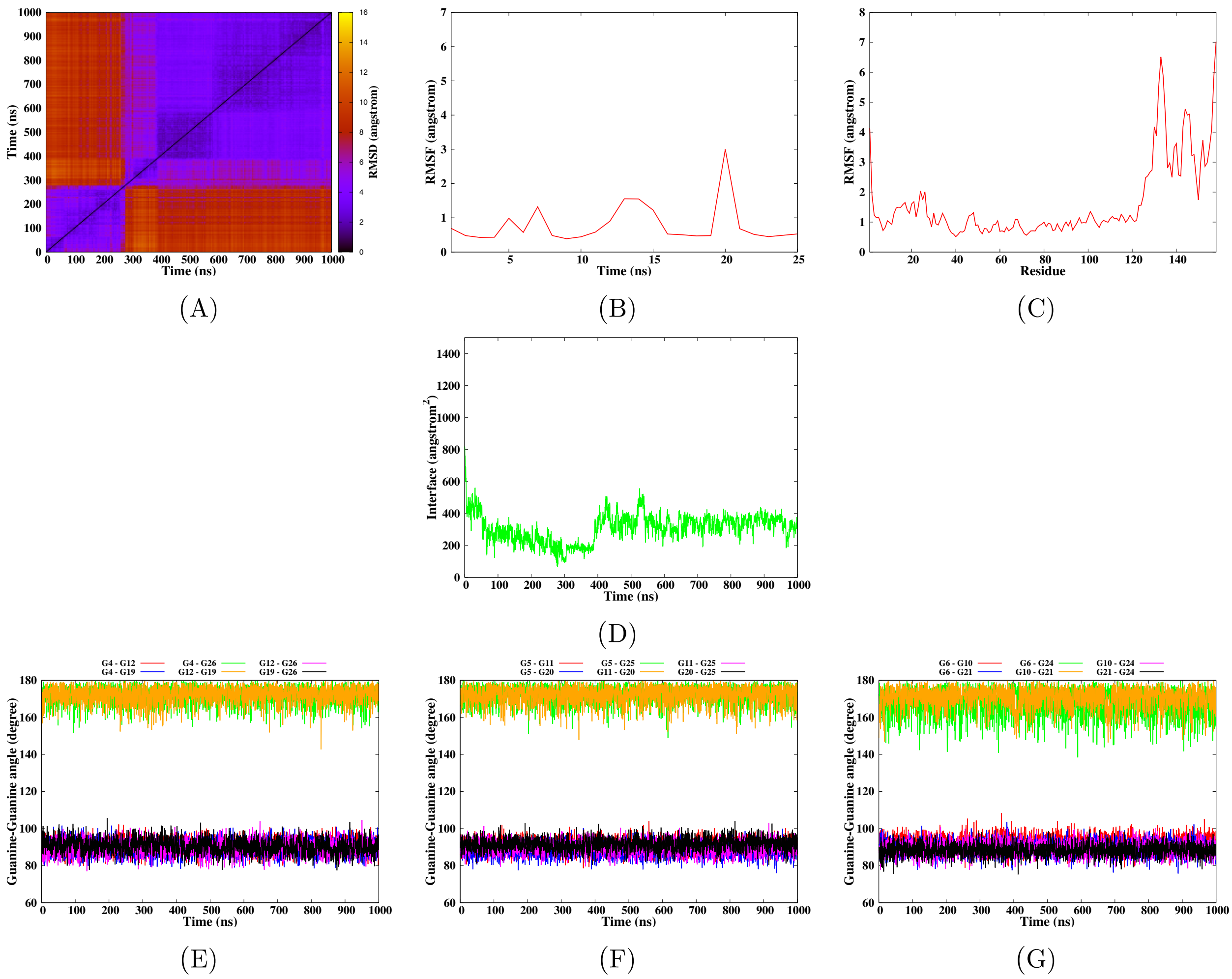


**Figure S34** – Simulation of the Bcl-2 G-quadruplex DNA in interaction with 2G10 according to the model 13-1, run 2. The convergence of the simulation is given by the RMSD-2D map of the DNA-Protein complex (A). The mobility of the DNA and protein residues is given by their root mean square fluctuation (B-C). Surface of the interaction interface between the protein and G-quadruplex (D). Finally the structural parameters of the G-quadruplex are given by the angles between the guanines for each tetrad (E-G).

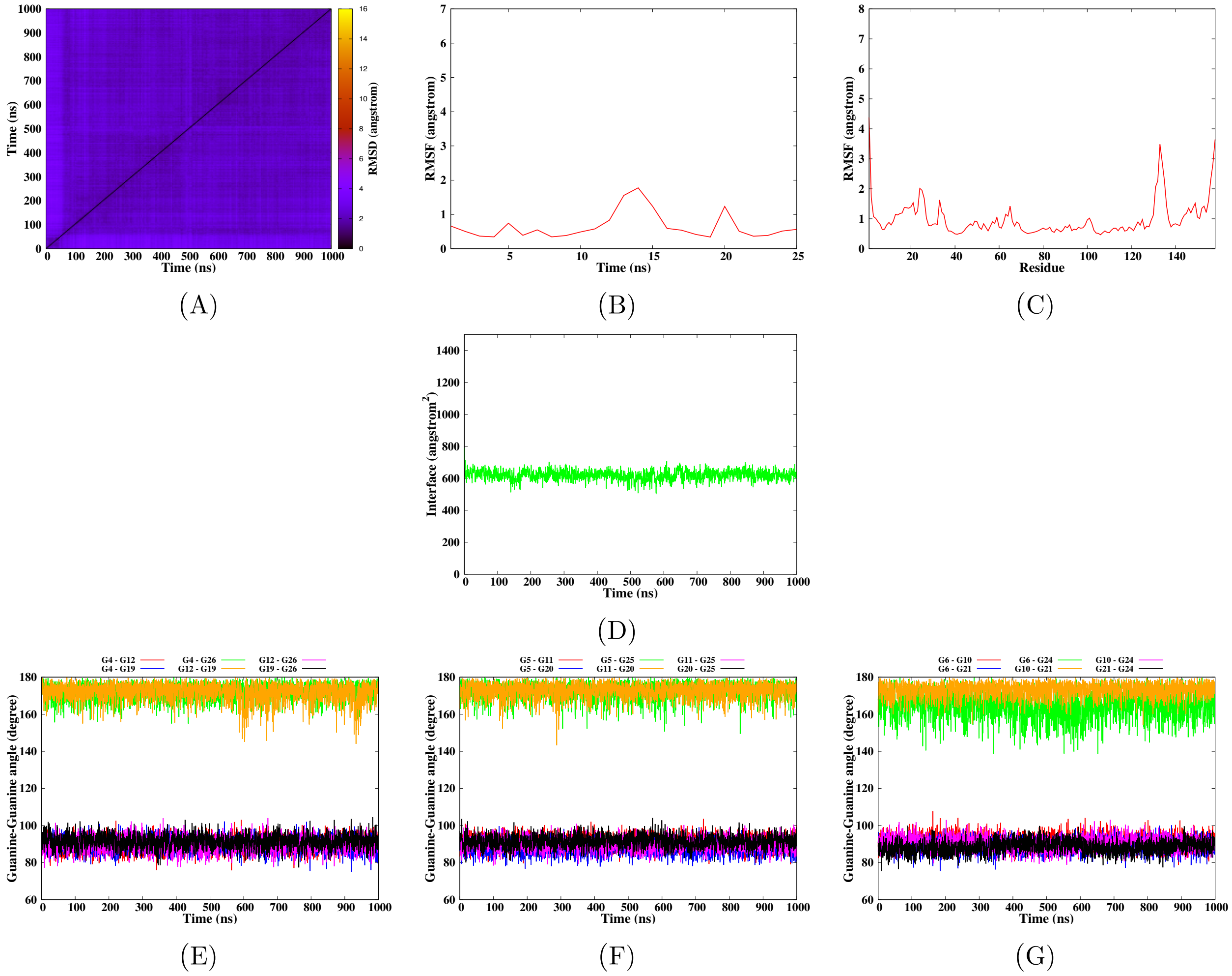


**Figure S35** – Simulation of the Bcl-2 G-quadruplex DNA in interaction with 2G10 according to the model 3-1, run 1. The convergence of the simulation is given by the RMSD-2D map of the DNA-Protein complex (A). The mobility of the DNA and protein residues is given by their root mean square fluctuation (B-C). Surface of the interaction interface between the protein and G-quadruplex (D). Finally the structural parameters of the G-quadruplex are given by the angles between the guanines for each tetrad (E-G).

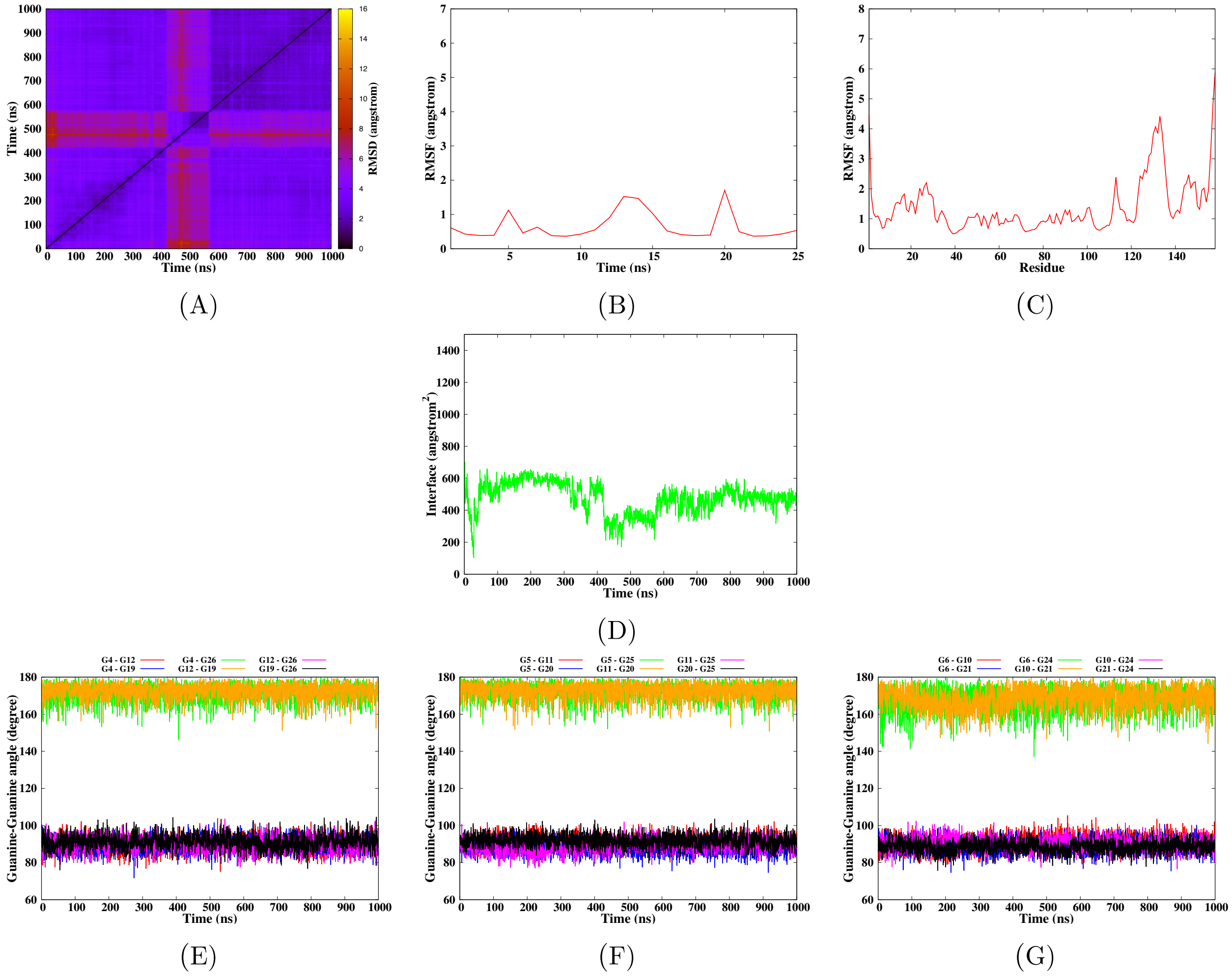


**Figure S36** – Simulation of the Bcl-2 G-quadruplex DNA in interaction with 2G10 according to the model 3-1, run 2. The convergence of the simulation is given by the RMSD-2D map of the DNA-Protein complex (A). The mobility of the DNA and protein residues is given by their root mean square fluctuation (B-C). Surface of the interaction interface between the protein and G-quadruplex (D). Finally the structural parameters of the G-quadruplex are given by the angles between the guanines for each tetrad (E-G).
